# A translational approach to measuring the neural systems underlying approach-avoidance conflict in humans

**DOI:** 10.1101/2022.07.27.501746

**Authors:** Camarin E. Rolle, Ken Amemori, Noriah Johnson, Marvin Yan, Trevor Caudle, Ana Havelka, James J. Gross, Amit Etkin

## Abstract

Approach-avoidance conflict (AAC) arises from decisions with embedded positive and negative outcomes, and appropriate management of these decisions is essential for adaptive functioning. However, translating key advances on AAC in non-human primates to tasks in humans has proven difficult in part due to the inherent limitations in existing human tasks in isolating relevant neural substrates of behavior. Here, we present, and validate, a novel task in humans (N= 38) of both sexes, derived from work in non-human primates utilizing primary reinforcers (shock/juice), and in doing so identify neural features specific to conflict, implementing a computational model of task behavior. We found that neural patterns of activation within the parietal, frontal, temporal and cingulate regions were associated with conflict-specific avoidance behavior. Importantly, a number of these regions were associated with trait anxiety, implicating a potential link between these neural regions and anxiety-driven avoidance behavior. This task platform may help advance both behavioral and biological research examining the neural patterning underlying approach-avoidance behavior in humans, providing an empirically oriented framework with which to translate between non-human primate and human work.

**Significance Statement:** The current paper describes and validates a novel task for studying Approach-Avoidance Conflict behavior that capitalizes on innate reinforcers and stringent thresholding, effectively bridging animal and human work in this field. The task is then used to identify the neural correlates of AAC behavior in humans, identifying patterns of activation linked to temporal variation in avoidance behavior specific to conflict. Further, a novel model is developed and applied to the behavioral data to more sensitively quantify response patterns using reinforcement learning, and neural patterns relating to these behavioral effects were identified. Finally, as anxiety is strongly associated in the animal and human literature with avoidance responding, we identified a subgroup of neural effects associated with trait anxiety.

## Introduction

It is well known that positive (i.e., desirable) outcomes generally drive approach behavior, while negative (i.e., undesirable) outcomes generally drive avoidance behavior [1]. However, in contexts in which outcomes are both positive *and* negative, behavior is more variable, and conflict is generally higher [1, 2]. This is termed “approach-avoidance conflict” (AAC)[3, 4] and we frequently face such conflict-laden decisions in our day-to-day interactions. In such situations, it is our idiosyncratic balance of “reward” and “punishment” sensitivities that ultimately determines how we act [1, 5, 6].

An imbalance between approaching and avoiding in such conflict decisions is thought to be a hallmark of a number of psychiatric disorders [7]. Imbalances favoring avoidance-motivated behavior, posited to underlie anxiety and neuroticism [8, 9], are symptomatic of anxiety-related disorders [10, 11]. In contrast, disproportionate approach-motivated behaviors, associated with greater impulsivity and extraversion [12], is thought to be a key feature of addiction [13], attention deficit hyperactivity disorder [14], and bipolar disorder [15]. Unfortunately, imbalances in decision-making can potentiate cyclic patterns of behavior leading to sustained, and even enhanced, disordered symptomatology [16]. The importance of understanding the mechanisms underlying variability in approach-avoidance conflict has motivated a rich body of work seeking to elucidate brain bases in non-human animals.

A large number of tasks in non-human animals and humans have been created to identify the predominant neural substrates driving AAC, and characterize the role of anxiety in driving avoidance behavior during conflict [17–20]. A critical step in building the bridge between basic neuroscience and clinical therapeutics is the effective translation of findings from the vast body of animal work to humans. However, the confounding sources of variability in task designs in humans limit the translatability of their findings to animal work [18]. There are two primary limitations of existing paradigms, the first of which is the lack of control for the subjective balance of embedded decision outcomes (e.g., the value of points versus negative pictures to a given person). If left unbalanced, there is a risk that the in-task behavior (and therefore simultaneous brain imaging) is influenced by how much the individual likes/dislikes the reward/punishment, contaminating the examination of the individual’s reward and punishment sensitivities.

The second limitation regards whether the task’s outcomes are evocative enough to drive sufficient conflict, which, if not, risks not motivating relevant approach-avoidance behavior.

To effectively study the neural bases of the biological drives motivating AAC in humans requires the removal of confounds and complexity that prevent definitive interpretation of the neural signals involved in the specific drives motivating behavior. In this way, it should be possible to begin to characterize the neural signatures most directly reflective of the drive towards reward and away from punishment, and how they interact to motivate behavior in AAC. The current paper describes a novel task for studying AAC behavior, and uses this task to identify the neural correlates of approach-avoidance conflict behavior. This task addresses a number of the limitations present in existing paradigms, and is validated to ensure its efficacy in doing so. As the task has been controlled and standardized to eliminate confounds (e.g., unbalanced outcomes) often encountered in the study of human behavior, it provides a strong translational platform to bridge primate and human work in studying approach and avoidance conflict. Grounded in the existant literature [18, 19, 21, 22], we hypothesize that the use of this task with simultaneous neural recordings will identify robust regions associated with approach-avoidance conflict, including the orbitofrontal, prefrontal, and cingulate cortex.

## Methods and Materials

### Participants

38 healthy adults (20 Female; Mean Age = 28.5 +/− 7.2 years) participated in this experiment. Written informed consent was obtained from each participant under institutional review board-approved protocols at Stanford University.

### Task Design

The task is modeled after a non-human primate study design for approach-avoidance conflict [23]. The task is comprised of four blocks in total, with each block parsed into two conditions: no-conflict (first-half) and conflict (second-half). During the task, two target stimuli are presented on either side of a computer screen (Figure 1). The participant is instructed that activating each stimuli (by pressing the keyboard button associated with it) will intermittedly result in juice being delivered. They are also informed that at some point within each block, one of the stimuli may intermittedly also result in shock, in addition to the juice. Each of the two targets is reinforced on an independent, variable ratio schedule, such that pressing both buttons as frequently as possible will result in the greatest amount of juice reward. Therefore, if a participant reduces their response rate to the “punished” stimuli (i.e., the stimuli associated with both juice and shock) once the shock is introduced during the block, they will receive less juice overall.

**Figure 1.**
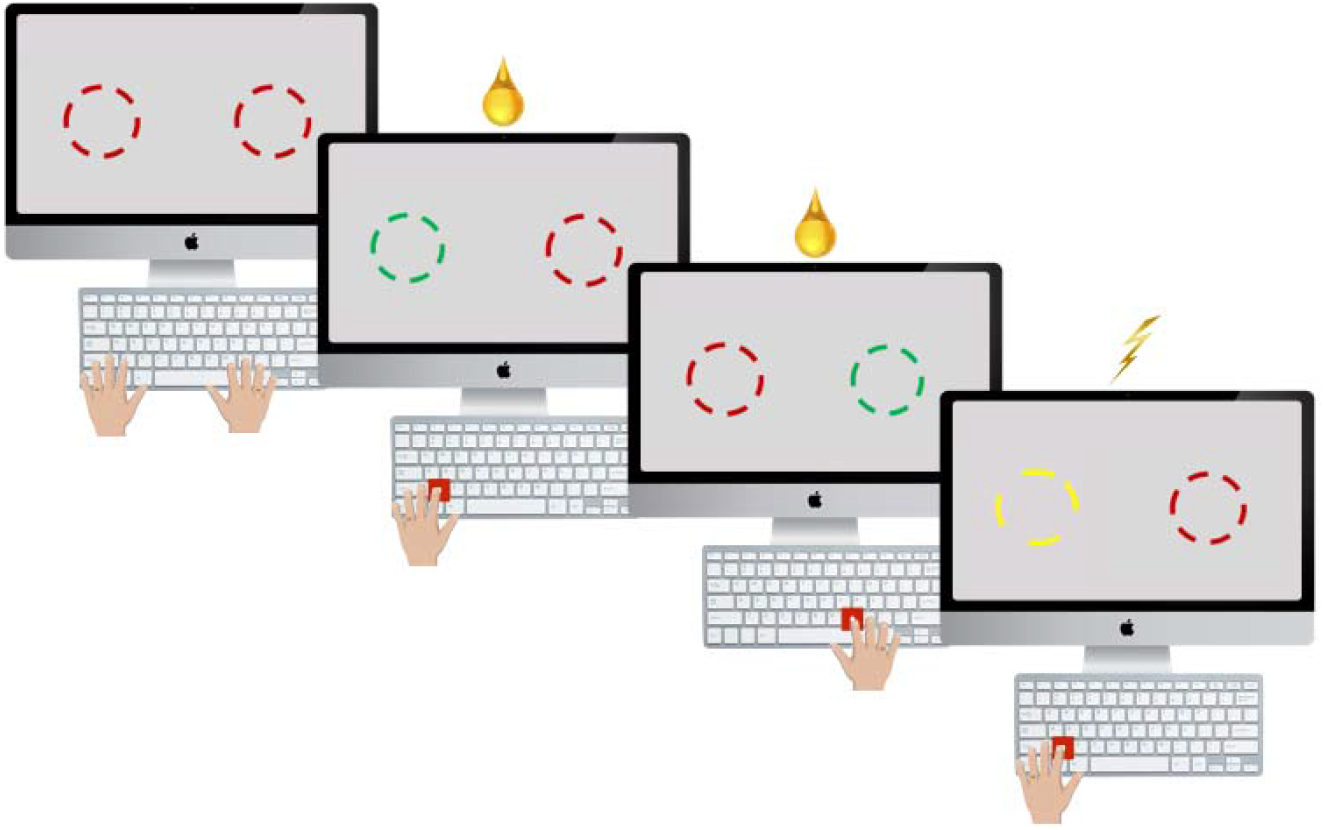
Task Design. Participants are instructed that left and right button pushes results in juice, and one of those button pushes (either left or right) will sometimes also results in shock. Participants are instructed that using both buttons will results in more juice than only using one.

**Figure 2.**
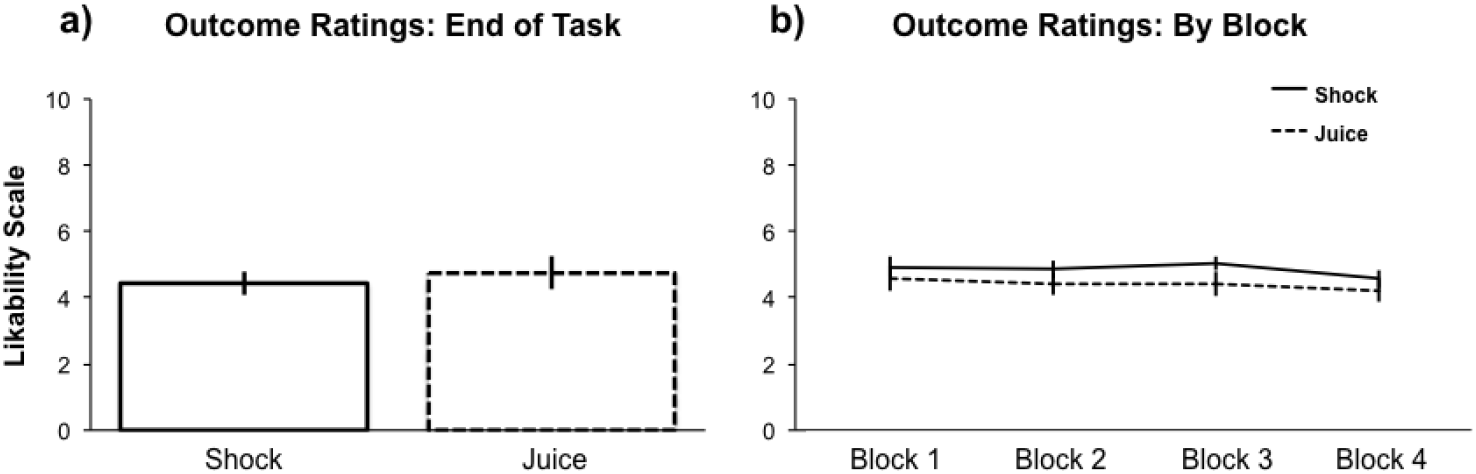
Thresholding Efficacy: a) Comparing likability between juice and shock in a post-task survey revealed equivalent, and opposite, preference ratings (T(39)=-.51, p=.61). b) Assessment of juice and shock “likability” across blocks revealed consistent reporting of juice and shock preference across blocks (ICC: Juice = .96, Shock=.85)

The shock is introduced halfway through a given block, with a jittered onset, and the stimuli it is associated with (left versus right target stimuli) is randomized across blocks so that one block is not predictive of the next one. Further task details and parameters can be found in the supplemental materials.

#### Measurement

As in the non-human primate study [23], approach was assessed by response bias, measured as the percent of responses, within a block, on the punished compared to non-punished side in the no-conflict condition (no shock administered) compared with the conflict condition (shock administered). Response bias was also extracted in 10-second bins to align with EEG estimates across time for the linear models.

#### Decision outcomes

In this task, juice was used as the reward, and shock was used as the punishment. The use of these innate reinforcers as the reward and punishment outcomes eliminated the confounding factor of conditioning, and reduced the subject-level variability in the outcome’s value seen with, for example, points and unpleasant pictures. Additionally, these particular reinforcers used in our task allowed for in-depth thresholding to ensure balanced and evocative in-task outcomes.

#### Shock Workup

To standardize the aversiveness of the punishment across participants, we embedded a shock workup in the procedure, in order to ensure that the in-task punishment resulted in equivalent discomfort across individuals. Details on this procedure can be found in the supplemental methods.

#### Juice Workup

Given humans’ ability to self-satiate (i.e., hydrate when thirsty), there may be attenuation of the hedonic value of a juice reward for humans as compared to primates. Therefore, we took a number of steps to ensure that juice had a substantial motivational value for each participant, including surveying juice preference, pre-testing fasting, and delivering oyster crakers. More details on the juice workup can be found in the supplemental materials.

#### In-Task Shock-Juice Equilibration

A number of in- and post-task surveys were administed to assess 1) The preference for each of the outcomes, 2) The stability of these preferences across blocks and 3) How matched the intensity of dislike for the shock was with the intensity of like for the juice. To assess the stability of this rating over blocks, we looked at agreement across block using Cronbach’s alpha intraclass correlation estimate [24]. Further details on these assessments can be found in the supplemental methods.

Following the workup, participants performed two in-task thresholding blocks, directly pairing juice and shock against each other, designed to ensure that the aversivesness of the shock was in fact calibrated to the appetitiveness of juice for each person. Specific details on these thresholding blocks are provided within the supplemental methods.

### Behavioral Analyses

#### Response Bias

To understand in-task behavior, and its stability across blocks within participants, we quantified individual behavior across the four blocks. We first looked at response bias to understand how the introduction of punishment was affecting responding by quantifying response bias. To assess the stability of this behavior over blocks, we looked at agreement across block using Cronbach’s alpha intraclass correlation estimate [24]. Response bias is quantified as the ratio of total responses that were “approach”: total responses on punished side / total responses. We expected response bias to be .5 during non-conflict trials as there was no shock being delivered, and therefore both right and left responses should be used equally. However we expected that during the conflict condition response bias would be less than .5, indicating a bias away from the punished response side.

Response bias was subdivided and averaged into 10-second bins for each subject, concatenated across blocks, to regress against spectral power in the linear mixed effects models outlined below.

#### Action Value

To more sensitively characterize choice patterns, we introduced a reinforcement learning computational model [25] to characterize the development of the biased choice pattern [26]. The model assigns an action value, *Q*(*t*), where *t* is the index of the trial for each action (i.e., right and left target selection). Q_Bias_ was ultimately quantified as Q_Non-Punished Side_ - Q_Punished Side_, representing the bias in action values associated with “avoid” responses compared to the “approach” responses. Therefore, we anticipate Q_Bias_ to increase as a function of avoidance following the introduction of punishment.

Q_Bias_ was subdivided and averaged into 10-second bins for each subject, concatenated across blocks, to regress against spectral power in the linear mixed effects models outlined below.

### EEG Data Acquisition

The EEG data was obtained using a 256-channel HydroCel Geodesic Sensor Net (HCGSN, Electrical Geodesics, Inc., Eugene, OR). Prior to data acquisition, the net was soaked in an electrolyte mixture of potassium chloride (KCl) with water (Electrical Geodesics, Inc. Eugene, OR) for 15 minutes. Each sensor was confirmed as being in contact with the participant’s scalp. Following net placement, electrode-scalp (resistive) impedances were measured to ensure that all electrode-scalp impedances were below 20 kΩ.

### EEG Data Processing

All EEG data analyses were performed in Matlab (R2014b, The Mathworks Inc., MA) using custom scripts built upon the preprocessing packages in EEGLAB [27] and ARTIST [28] toolboxes. Details of data clearning can be found in the supplemental methods. Cleaned data were binned into 10-second time bins, concatenated across blocks. For each subject, an imaging kernel that maps from the channel space EEG to the source space current density was then estimated by the minimum norm estimation approach with depth weighting and regularization, and rotating dipoles at 3003 vertices were generated on the cortical surface.

### EEG Spectral Analyses

All spectral analyses were computed at the vertex level using 3003 vertices in MNI template space. Data underwent morlet wavelet time frequency transformation, and then the power was averaged within the four canonical frequencies: theta (4.5-7.5Hz), alpha (8-12Hz), beta (12.5-30Hz), and gamma (31-50Hz) frequency bands. Finally, power estimates for each individual were normalized across the 3003 vertices, using a Z-transformation statistic [29].

### Statistical Analyses

For each individual, behavioral and EEG data were subdivided and averaged into 10-second bins of data, concatenated across all blocks. In doing so, bin-to-bin variation in behavior (response bias and Q) was used as a continuous regressor against bin-level band power.

Linear mixed-effects models were applied to each vertex band power estimate [30]. Analyses were conducted using the NLME package in R [31]. The modeled effects included a random intercept, and fixed effect of task condition (conflict, no-conflict) and trial number (10-second bins of data). These effects were modeled by an interaction with the source-level spectral estimates (EEG) examined to identify which EEG estimates were associated with behavior. The primary term of interest is the EEG x condition interaction, which assessed bin-to-bin variability in response bias (% approach) and Q_Bias_ as a function of EEG spectral band moderators. EEG spectral features that varied as a function of condition were assessed by a significant condition main effect using the identical linear mixed effects model, with fixed effect of task condition (conflict, noconflict) and trial number (10-second bins of data), and source-level spectral estimates (EEG). Voxels found to associate with behavior differentially depending on task condition were then grouped into clusters (details of which can be found the supplemental methods).

All p-values were FDR-corrected for multiple comparisons across all vertices and frequency bands to control for type I errors.

To further understand the potential clinical significance of behavior-moderating spectral features, we ran Spearman’s correlations of the FDR-significant EEG results with the trait component of the State-Trait Anxiety Inventory [32] (correcting for multiple corrections across all correlations run) across the entire participant sample.

## Results

### Task Validation

#### Shock and Juice “Likability” Ratings

In-task ratings revealed that the task elicited motivationally-evocative, balanced outcomes across participants. The specific results to all assessments leading to this conclusion can be found in the supplemental results.

#### In-Task Shock-Juice Equilibration

Results suggested a variable but effective thresholding which led participants, on average, to approach juice reward despite the associated shock punishment approximately 60% of the time. Further quantification of Shock-Juice equilibration can be found in the supplemental results.

#### In-Task Behavior

Participants averaged 152.62 +/− 53.97 responses/minute, with each block taking an average of 6.14 +/− 2.23 minutes. In-task % approach response of total responses (*see* Figure 3a) was 50.56%+/− 12.75% for no-conflict conditions, and 23.64%+/−15.89% for conflict conditions (T(37)=-8.44, p<.001), with an average of 26.92%+/− 19.64% increase in response bias away from punished side in conflict conditions compared to non-conflict conditions. Agreement in response bias across conflict conditions was .92, and across no-conflict conditions was .57 (*see* Figure 3b). Q_Bias_, reflecting the action value of the response associated with non-punished side compared to the punished side, was significantly different between conflict and nonconflict conditions (T(37)=-6.16, p<.001; Figure S1a). Agreement in response bias across conflict conditions was .95, and across no-conflict conditions was .53 (*see* Figure S1b).

**Figure 3.**
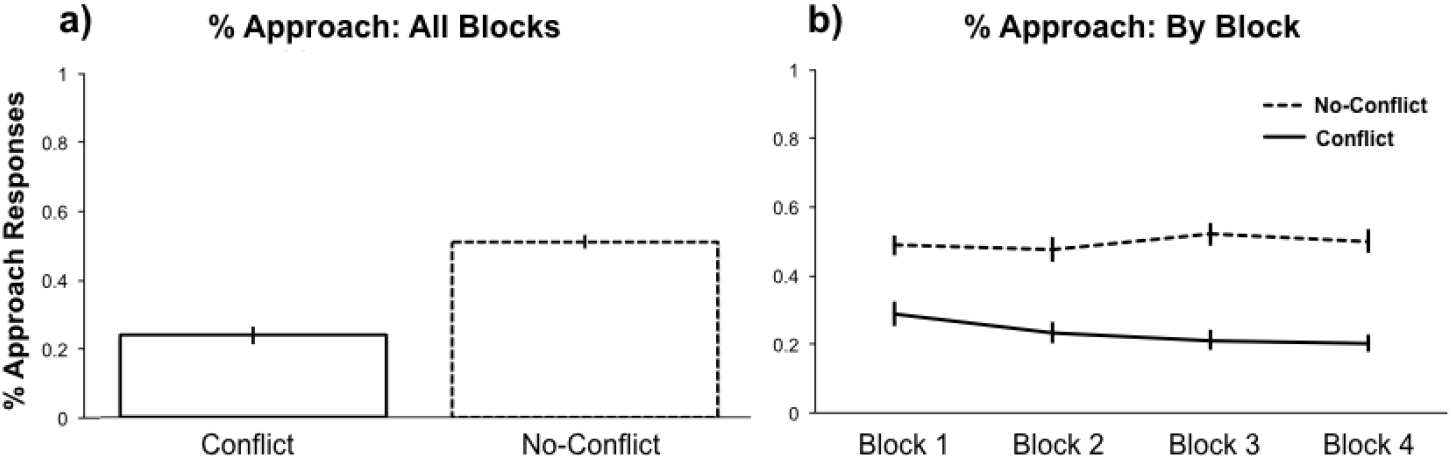
Assessment of in-task response Bias: a) % approach responses was 50.56% +/− 12.75% for no-conflict conditions, and 23.64% +/−25.89% for conflict conditions (T(37)=-8.44, p<.001), with an average of 26.92% +/− 19.64% increase in response bias away from punished side in conflcit conditions compared to no-conflict condidtions. b) Stability of in-task behavior across blocks was assessed by ICC (Conflict = .92, No-Conflict = .57).

### EEG

#### Main Effects of Condition

After correcting for multiple comparisons across all 12012 (3003 vertices, 4 spectral bands) EEG features assessed, 175 distinct clusters were found to statistically differ by task condition, evidenced by a main effect of condition (see Table S1; Figure 4). These results showed a number of clusters with peak voxel activation in the following regions to have lower power during conflict compared to no-conflict conditions: right anterior prefrontal cortex (aPFC), bilateral associative visual cortex, bilateral cingulate cortex, bilateral dorsal lateral prefrontal cortex (dlPFC), bilateral frontal eye fields (FEF), right fusiform gyrus, left inferior temporal gyrus (ITG), left middle temporal gyrus (MTG), left orbitofrontal cortex, right premotor cortex, and right primary auditory cortex. These results showed a number of clusters in the following regions to have greater power during conflict compared to no-conflict conditions: bilateral angular gyrus, left anterior prefrontal cortex, bilateral associative visual cortex, bilateral cingulate cortex, bilateral dlPFC, right FEF, right ITG, bilateral MTG, bilateral orbitofrontal cortex, bilateral premotor, right somatosensory, bilateral superior temporal gryus, bilateral supramarginal gyrus, bilateral superior parietal, and right brocas area. While the majority of effects were unspecific to a particular frequency band, cingulate activation was most pronounced within the theta band, and lateral prefrontal activation most pronounced in theta and alpha bands.

**Figure 4.**
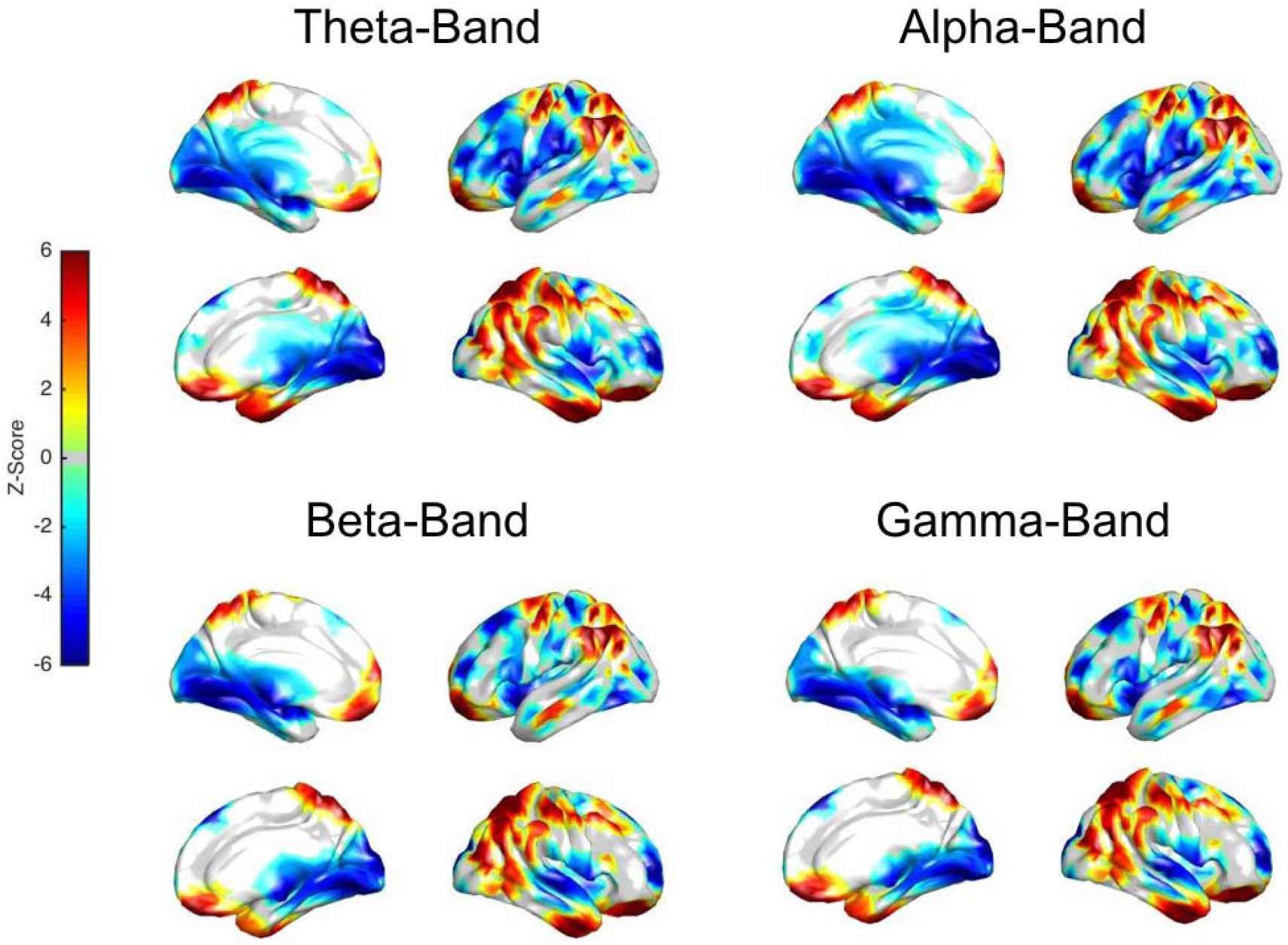
Significant condition main effects. Cortical representation of z-values for EEG features significantly differing by condition. Positive z-scores denote reduced power during conflict, whereas negative z-scores denote enhanced power during conflict.

#### Condition x Avoidance Interaction Effects

After FDR-correcting for multiple comparisons across all 12012 (3003 vertices, 4 spectral bands) EEG features assessed, 167 distinct clusters were found to associate with bin-to-bin variability in in-task avoidance behavior differentially by task condition, as measured by response bias (see Table S2; Figure 5a). The majority of these clusters were found within frontal and parietal cortices, with the rest falling within the visual, somatosensory/motor, temporal, or cingulate cortices. While the majority of effects were global across frequency bands, cingulate activation was most pronounced within the alpha and theta bands.

**Figure 5.**
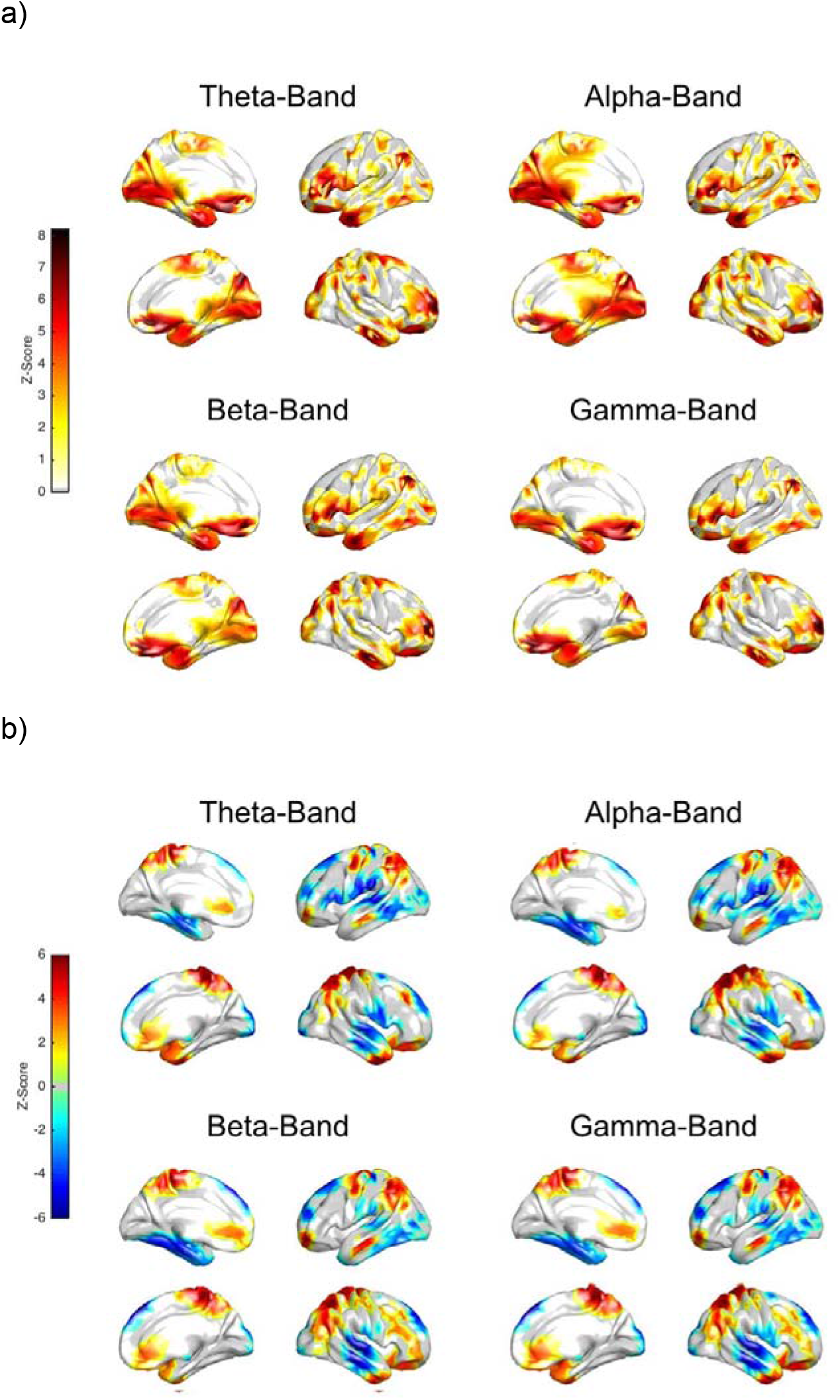
Significant moderators of task avoidance (Response Bias). a) Cortical representation of z-values for significant ROI moderators of task avoidance. b) Cortical representation of z-values (color coded to represent directionality) for significant ROI predictors of conflict-specific task avoidance are represented. Positive z-scores denote a negative association within the conflict condition between EEG band power and % avoidance, whereas negative z-scores denote a positive association.

We next sought to understand which of these significant EEG features were driven by behavior-EEG associations specific to the conflict condition (Figure 5b). We ran mixed models for the two task conditions separately, further constraining analyses to only those EEG features identified as significant in the full analysis using a p-value threshold of 0.05. Greater power was associated with less % avoidance behavior during conflict trials predominantly within cingulate, frontal and parietal cortices, whereas greater power was associated with more % avoidance during conflict trials within the temporal and sensory cortices (i.e., motor, visual, auditory). While the majority of effects were again global across frequency bands, cingulate, lateral prefrontal, and medial somatosensory and temporal activation was most pronounced within the alpha and theta bands.

Q_Bias_ showed a similar pattern of results as response bias (see Table S3; Figure 6a). Greater power predicted less Q_Bias_ behavior during conflict trials within frontal and visual areas, whereas greater power predicted greater QBias behavior during conflict trials within temporal and motor areas. While the frontal and visual effects were found across frequency bands, the temporal effects were isolated within the alpha and theta bands.

**Figure 6.**
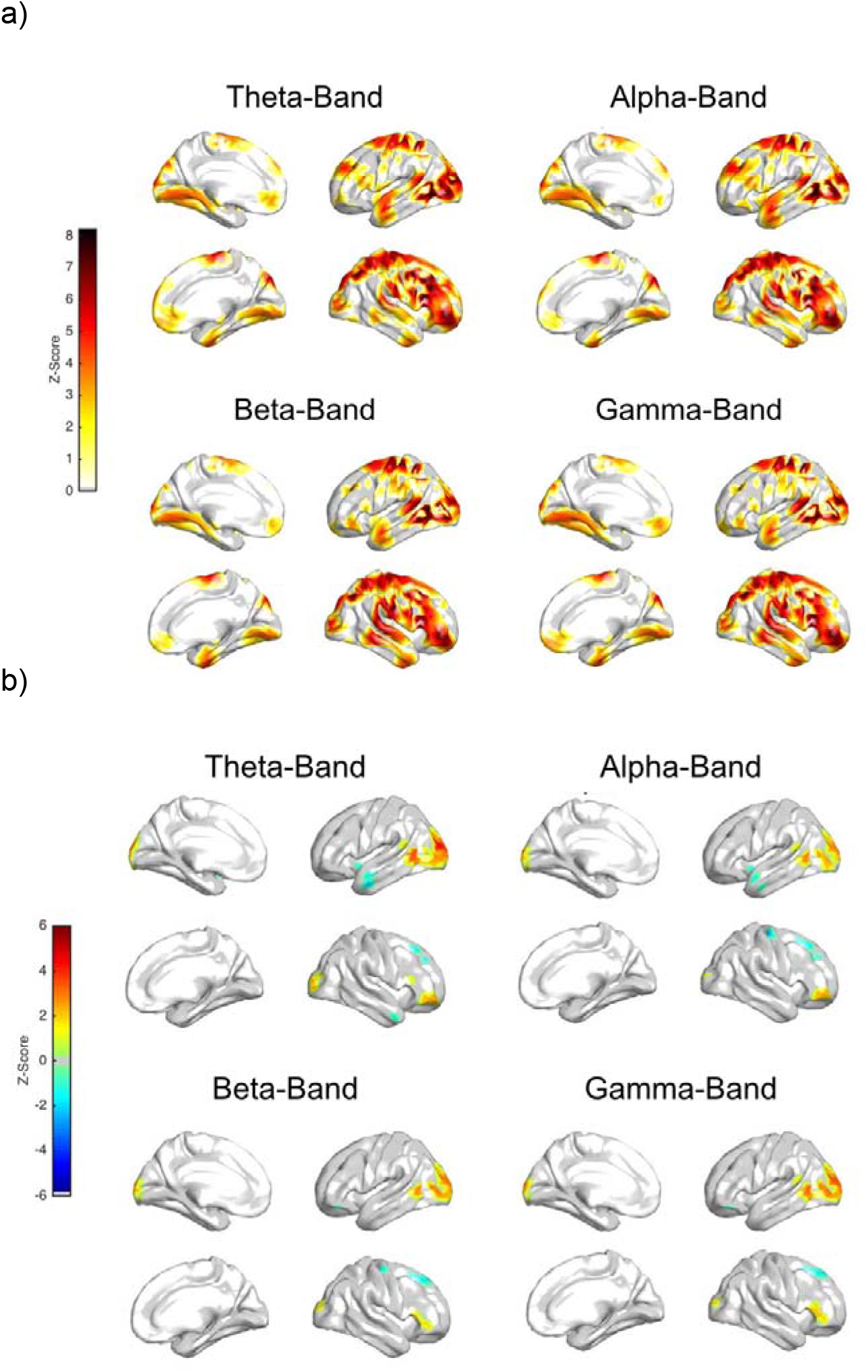
Significant moderators of task avoidance (Q_Bias_). a) Cortical representation of z-values for significant ROI moderators of task avoidance. b) Cortical representation of z-values (color coded to represent directionality) for significant ROI predictors of conflictspecific task avoidance are represented. Positive z-scores denote a negative association within the conflict condition between EEG band power and % avoidance, whereas negative z-scores denote a positive association.

### STAI-T Correlations

After correcting for multiple comparisons across all 12012 (3003 vertices, 4 spectral bands) EEG features assessed, a large proportion of significant voxels identified in the LME analyses were found to associate with STAI-T (see Figure 7). The majority of surviving voxels negatively associated with trait anxiety were within the frontal, cingulate and parietal regions, whereas those positively associated with anxiety were predominantly temporal. While the temporal and prefrontal effects were shown across all bands, the cingulate was most pronounced within the alpha and theta bands.

**Figure 7.**
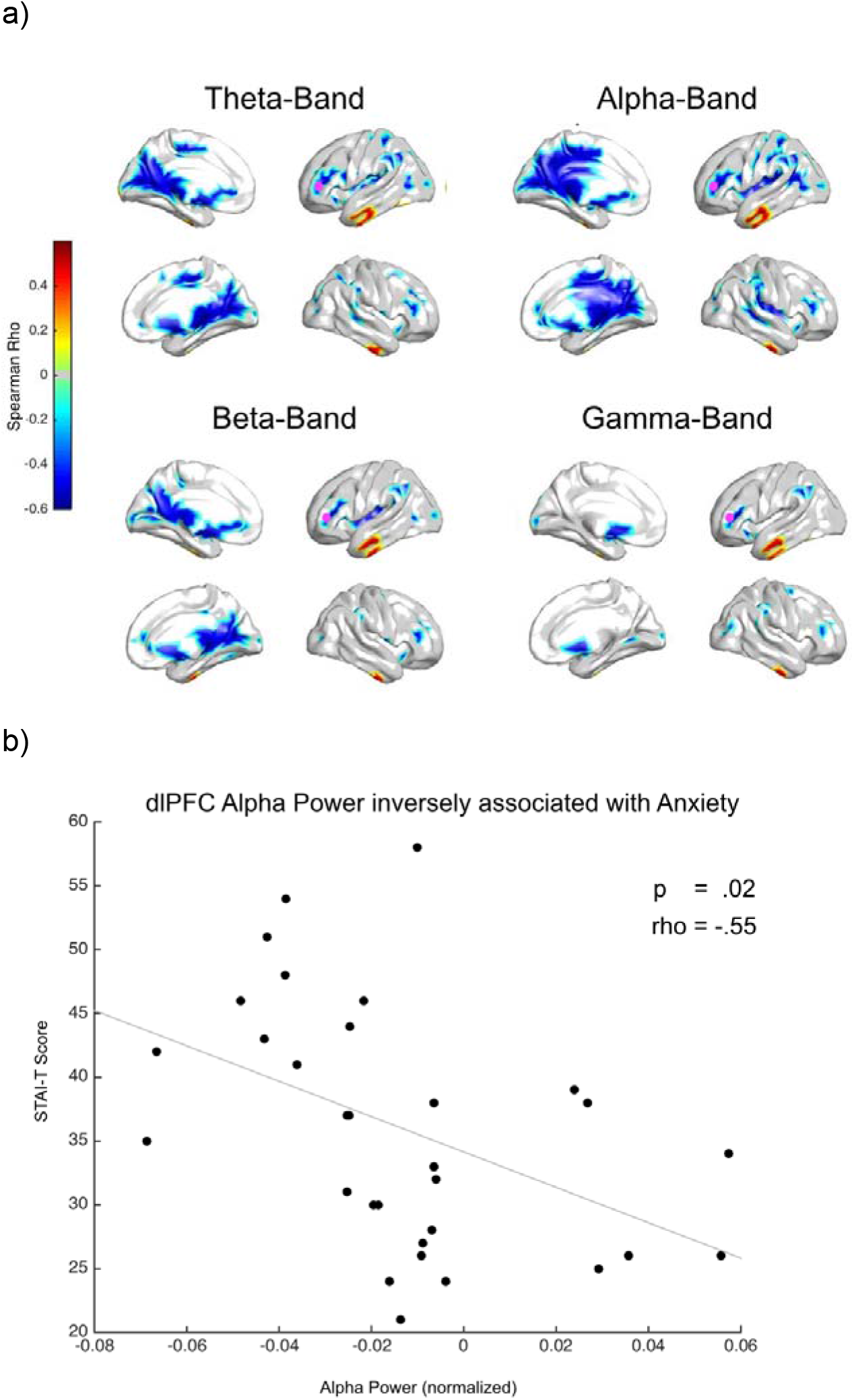
Vertices significantly associated with anxiety. a) Cortical representation of rho-values for vertices significantly associated with anxiety, as measured by STAI-T. Visualization is masked by vertices whose band power was significantly associated with conflict-specific response bias in the LME analyses. b) Example scatter plot demonstrating negative association between dorsal lateral prefrontal cortex (dlPFC) alpha power and trait anxiety. Vertex plotted in (b) is highlighted in magenta in (a).

### STAI-T Correlations

After correcting for multiple comparisons across all 12012 (3003 vertices, 4 spectral bands), no associations between Q_Bias_ and EEG power survived.

## Discussion

The current study proposes a novel task, fashioned after non-human primate work, and uniquely positioned to isolate relevant neural circuitry of approach-avoidance behavior by its layered thresholding procedures, and use of primary reinforcers. The validation of the proposed paradigm (further discussion in the supplemental materials) suggests that 1) the thresholding effectively produces balanced and evocative reward and punishment, and 2) participant behavior elicited in this task is relatively stable. As such, the current task provides a powerful translational bridge to animal work. Therefore, the EEG features found in this study to associate with bin-to-bin variability in behavior, differentially by task condition, closely reflect those most relevant to conflict-specific approach- and avoidance-drives.

Importantly, the task then identifies a number of cortical features whose band power (1) dissociates conflict from non-conflict conditions within the task, (2) differentiates task condition in its association with avoidance behavior and (3) associates with trait anxiety. The study does this through the derivation of two behavioral metrics, one quantified from response patterns, and the second from a reinforcement learning model aimed to enhance sensitivity by quantifying bin-to-bin variability in behavior through learning rate and cost-benefit ratio estimates.

### EEG regions Associated with Avoidance Behavior during Conflict

Using this task, we found a number of spectral features that associated with avoidance behavior differentially between conflict and no-conflict conditions. These features were shared across all spectral bands, and fell predominantly within the frontal, parietal, temporal and cingulate cortices.

The features predictive of avoidance behavior in conflict trials were predominantly found in the bilateral orbitofrontal, anterior and dorsolateral prefrontal cortex, superior, inferior and middle temporal gyrus, and cingulate. Interestingly, the majority of these features were not specific to a single frequency band, and generally activated within most, if not all, of the four spectral bands. Specifically, greater frontal and cingulate spectral power associated with less avoidance behavior during conflict trials, whereas less temporal and motor power associated with greater avoidance during conflict. Our findings are well aligned with much of the fMRI literature studying the neural correlates of approach-avoidance conflict.

Previous work has implicated the orbitofrontal, prefrontal, and cingulate cortex to be particularly prominent in approach-avoidance conflict [18, 19, 21, 22]. The orbitofrontal cortex, known to reflect reward sensitivity [33–36] and to play a role in regulating limbic regions during emotion processing [37–41], potentially through its anatomical connectivity with the limbic system [41], has shown greater activation during conflict trials in both humans and non-human primates [18, 19, 22, 42]. The anterior mid-frontal gyrus, making up the anterior part of the dorsal-lateral PFC, is a part of the salience network, most commonly implicated in attentional processes, particularly conflict [43]. The dlPFC is arguably the most commonly implicated cortical region in approach-avoidance conflict behavior [17, 18, 44, 45], thought to act on these processes through its functional role in goal motivation. The cingulate, commonly associated with signaling conflict [46, 47], particularly in the context of emotion [39–41], is thought to engage the dlPFC for regulation of limbic processes [47]. The cingulate has been shown to be commonly implicated in approach-avoidance conflict [18], with microstimulation to this area in non-human primates leading to greater avoidance [48].

The temporal cortex, though known to be highly innervated with connections to the amygdala [49] and involved in emotion regulation [50], has been largely neglected in the field of approach-avoidance conflict. However, we find the temporal cortex to positively associate with avoidance during conflict, such that greater temporal cortex power aligned with greater avoidance behavior. An existent theory suggests that the amygdala primes representations of emotional stimuli in temporal cortex, causing an augmented focus of attention on emotional content compared to non-emotional content [51]. We therefore hypothesize that it is in its connectivity with the amygdala that we see limbic-like patterns of activity in this cortex, such that greater power in the temporal cortices is associated with greater avoidance.

Our primary output from the computational model, Q_Bias_, replicated the majority of those EEG features found to associate with response bias during conflict tasks. We adopted the reinforcement learning model to characterize the change in the action value. The action values are the internal variables estimated by the adaptive change in value judgments, and represent the predictive value of the expected outcome for each trial. We introduced the action value derived from the reinforcement learning model so that we could capture the internal variables necessary and critical to derive the adaptive change in the valuation.

### Interpretation of Broadband Spectral Signal

The majority of effects observed in the current study were not specific to a single frequency band. As our task was blocked and absent of a trial structure, measuring state rather than evoked effects, it is unsurprising that many of the effects were broadband (Stephen et al, 2019; Mukamel et al 2014; Winawer et al 2013). It has been shown that generalized broadband activation, and not band-specific oscillations, reflects population-level neuronal spiking (Kreiman et al., 2006; Manning, Jacobs, Fried, & Kahana, 2009; Miller et al., 2007; Miller 2010; Kupers et al 2019). Fluctuations of electrical potential are generally assumed to be the sum of more or less narrow-band oscillatory activities in distinct frequency bands (Onton and Makeig, 2009), and broadband electrophysiological signal most closely correlates with functional Magnetic Resonance Imaging (fMRI) blood-oxygen level dependent (BOLD) signal (Winawer et al 2013). From this, we could infer that the findings shared across the narrow frequency bands, predominantly parietal, could be a less process specific, and more related to general activation (Wen and Liu, 2016; Pan et al 2011).

While the majority of effects were global across frequency bands, cingulate, lateral prefrontal, and medial somatosensory and temporal activation was most pronounced within the narrow low-frequency (i.e., alpha and theta) bands, perhaps suggesting their involvement in more process-focused aspects of conflict throughout the task blocks. This is particularly interesting because theta and alpha are thought to be causally relevant to affective processing (e.g., Zhao et al 2018), conflict processing (e.g., Jian et al 2018; Chinn et al 2018;), and generalized anxiety (Dadashi et al 2015; Moore 2000), so much so that modulation of these particular oscillatory bands are foundational mechanisms to some therapeutic treatments for anxiety disorders (Moore 2000). However, as the literature on the relevance of narrow-band spectral power to cognitive processing is vastly heterogeneous, the relationship between alpha- and theta-band oscillations and approach-avoidance conflict would need to be validated through more directed testing.

### Relationships Between Associated EEG Features and Anxiety

Anxiety is heavily implicated in approach-avoidance conflict behavior in both human and animal literature. The animal literature has a wealth of data suggesting the causal role of anxiety in avoidance, with anxiolytic administration consistently decreasing avoidance behavior [52, 53]. Human work has corroborated the relevance of anxiety to conflict behavior, with heightened anxiety consistently associating with greater avoidance responding in approach-avoidance conflict [17, 18, 20, 21].

We found negative correlations between trait anxiety as measured by the STAI-T, and several of the spectral features found to negatively predict avoidance behavior, including frontal, cingulate and parietal power, most predominantly seen in the lower-frequency narrow bands (i.e., alpha and theta). Further, we found positive correlations between trait anxiety and temporal power. As avoidance has been shown to be elevated in those with anxiety [17, 18, 20, 21], these findings are in line with our findings that frontal, cingulate and parietal regions negatively associate, and temporal power positively, with avoidance behavior specific to conflict. This points to the relevance these vertices to anxiety, and could either suggest that heightened anxiety accompanies reduced power in these regions, which accompanies greater avoidance behavior, or potentially point to anxiety as a mechanism through which these regions’ activation impact conflict behavior. Further testing to causally elucidate the role of both anxiety and these cortical regions in approach-avoidance behavior are required to more comprehensively understand their relationship.

## Limitations

Using a novel and validated task (the limitations of which can be found in the supplemental materials), we provided initial evidence regarding the brain bases of AAC. However, limitations should be noted. One limitation is that the current work provides little insight into the network-level underpinnings of approach-avoidance conflict, and would therefore benefit from connectivity work aimed to better understand the regional interactions accompanying various behavioral patterns. The current study was in no way causal, and therefore further work interrogating the role of these regions in approach-avoidance conflict is required to more definitively understand the relevance of these regions to behavior. Finally, the EEG was acquired using a high-density 256-channel cap which is known to improve inverse solution accuracy [54], source localization has known limitations given its inherent assumptions and its inability to localize depth sources beyond the cortical surface [55]. However, the minimum norm solution we apply has been shown to be optimally suited for the resolution and accuracy limits of EEG measurement [56].

## Conclusions and Future Directions

The current paper proposes and validates a novel task for studying approach-avoidance conflict in humans with translational promise to animal work. Using this task, we identify a number of key spectral features that significantly predicted conflict behavior, with a subset of these features associated with anxiety. Moving forward, we emphasize the importance of controlling for all aspects of the aforementioned experimental components in designing a study, and importantly, assessing the efficacy of doing so for the inclusion of such information in subsequent analyses. Given the inherent complexities and variability in human studies, further work in the field needs to be done to ensure that such a standard is held for all work studying psychological concepts in this population.

## Acknowledgements

The study was funded by NIH grant DP1 MH116506. Rolle, C was additionally funded by NSF GRFP grant (No. 2016180976).

## References

[1] P.J. Corr, Approach and Avoidance Behaviour: Multiple Systems and their Interactions, Emotion Review 5(3) (2013) 285–290.

[2] S. Roth, L.J. Cohen, Approach, avoidance, and coping with stress, The American psychologist 41(7) (1986) 813–9.

[3] R.A. Champion, Motivational effects in approach-avoidance conflict, Psychological review 68 (1961) 354–8.

[4] A.J. Elliot, T.M. Thrash, Approach-avoidance motivation in personality: approach and avoidance temperaments and goals, Journal of personality and social psychology 82(5) (2002) 804–18.

[5] P.J. Corr, N. McNaughton, Neuroscience and approach/avoidance personality traits: a two stage (valuation-motivation) approach, Neuroscience and biobehavioral reviews 36(10) (2012) 2339–54.

[6] D. Mobbs, J.L. Marchant, D. Hassabis, B. Seymour, G. Tan, M. Gray, P. Petrovic, R.J. Dolan, C.D. Frith, From threat to fear: the neural organization of defensive fear systems in humans, The Journal of neuroscience: the official journal of the Society for Neuroscience 29(39) (2009) 12236–43.

[7] M.B. Stein, M.P. Paulus, Imbalance of approach and avoidance: the yin and yang of anxiety disorders, Biological psychiatry 66(12) (2009) 1072–4.

[8] P.J. Corr, Reinforcement sensitivity theory and personality, Neuroscience and biobehavioral reviews 28(3) (2004) 317–32.

[9] J.A.N.M. Gray, Fundamentals of the septo-hippocampal system, in: O.U. Press (Ed.), The Neuropsychology of Anxiety: An Enquiry into the Functions of Septo-hippocampal System, Oxford, 2000, pp. 204–232.

[10] P. Muris, H. Merckelbach, H. Schmidt, B.B. Gadet, N. Bogie, Anxiety and depression as correlates of self-reported behavioural inhibition in normal adolescents, Behaviour research and therapy 39(9) (2001) 1051–61.

[11] O. Agcaoglu, T.W. Wilson, Y.P. Wang, J. Stephen, V.D. Calhoun, Resting state connectivity differences in eyes open versus eyes closed conditions, Human brain mapping 40(8) (2019) 2488–2498.

[12] J.A. Gray, The neuropsychology of temperament.” Explorations in temperament, Springer US (1991) 105–128.

[13] P. Bijttebier, I. Beck, L. Claes, W. Vandereycken, Gray’s Reinforcement Sensitivity Theory as a framework for research on personality-psychopathology associations, Clinical psychology review 29(5) (2009) 421–30.

[14] J.T. Mitchell, & Nelson-Gray, R. O., Attention-deficit/hyperactivity disorder symptoms in adults: relationship to Gray’s behavioral approach system, Personality and Individual Differences 40(4) (2006) 749–760.

[15] R.A. Chandler, J. Wakeley, G.M. Goodwin, R.D. Rogers, Altered risk-aversion and risk-seeking behavior in bipolar disorder, Biological psychiatry 66(9) (2009) 840–6.

[16] K. Mogg, & Bradley, B. P. (Attentional bias in generalized anxiety disorder versus depressive disorder., Cognitive therapy and research 29(1) (2005) 29–45.

[17] R.L. Aupperle, M.P. Paulus, Neural systems underlying approach and avoidance in anxiety disorders, Dialogues in clinical neuroscience 12(4) (2010) 517–31.

[18] N. Kirlic, J. Young, R.L. Aupperle, Animal to human translational paradigms relevant for approach avoidance conflict decision making, Behaviour research and therapy 96 (2017) 14–29.

[19] M.W. Schlund, A.T. Brewer, S.K. Magee, D.M. Richman, S. Solomon, M. Ludlum, S. Dymond, The tipping point: Value differences and parallel dorsal-ventral frontal circuits gating human approach-avoidance behavior, NeuroImage 136 (2016) 94–105.

[20] J. Sheynin, K.D. Beck, R.J. Servatius, C.E. Myers, Acquisition and extinction of human avoidance behavior: attenuating effect of safety signals and associations with anxiety vulnerabilities, Frontiers in behavioral neuroscience 8 (2014) 323.

[21] R.L. Aupperle, S. Sullivan, A.J. Melrose, M.P. Paulus, M.B. Stein, A reverse translational approach to quantify approach-avoidance conflict in humans, Behavioural brain research 225(2) (2011) 455–63.

[22] D. Talmi, P. Dayan, S.J. Kiebel, C.D. Frith, R.J. Dolan, How humans integrate the prospects of pain and reward during choice, The Journal of neuroscience: the official journal of the Society for Neuroscience 29(46) (2009) 14617–26.

[23] H.F. Clarke, N.K. Horst, A.C. Roberts, Regional inactivations of primate ventral prefrontal cortex reveal two distinct mechanisms underlying negative bias in decision making, Proceedings of the National Academy of Sciences of the United States of America 112(13) (2015) 4176–81.

[24] K.O. McGraw, S.P. Wong, Forming Inferences About Some Intraclass Correlation Coefficient, Psychological Methods 1(1) (1996) 30–46.

[25] C. Watkins, & Dayan, P., Q-learning, Machine Learning 8(3) (1992) 279–292.

[26] K. Katahira, The relation between reinforcement learning parameters and the influence of reinforcement history on choice behavior Journal of Mathematical Psychology 66 (2015) 59–69.

[27] A. Delorme, S. Makeig, EEGLAB: an open source toolbox for analysis of single-trial EEG dynamics including independent component analysis, Journal of neuroscience methods 134(1) (2004) 9–21.

[28] W. Wu, C.J. Keller, N.C. Rogasch, P. Longwell, E. Shpigel, C.E. Rolle, A. Etkin, ARTIST: A fully automated artifact rejection algorithm for single-pulse TMS-EEG data, Human brain mapping 39(4) (2018) 1607–1625.

[29] S. Tong, N.V. Thakor, Quantitative EEG analysis methods and clinical applications, Artech House2009.

[30] M.L. Lindstrom, D.M. Bates, Nonlinear mixed effects models for repeated measures data, Biometrics 46(3) (1990) 673–87.

[31] J. Pinheiro, D. Bates, S. DebRoy, D. Sarkar, Linear and nonlinear mixed effects models, R package version, 2014.

[32] C.D. Spielberger, R.L. Gorsuch, R.E. Lushene, Manual for the State-Trait Anxiety Inventory (SelfEvaluation Questionnaire), Consulting Psychologists Press, Palo Alto, CA, 1970.

[33] J.P. O’Doherty, Lights, camembert, action! The role of human orbitofrontal cortex in encoding stimuli, rewards, and choices, Annals of the New York Academy of Sciences 1121 (2007) 254–72.

[34] R.A. Wise, Dopamine and reward: the anhedonia hypothesis 30 years on, Neurotoxicity research 14(2-3) (2008) 169–83.

[35] E.T. Rolls, F. Grabenhorst, The orbitofrontal cortex and beyond: from affect to decision-making, Progress in neurobiology 86(3) (2008) 216–44.

[36] J.J. Simon, S. Walther, C.J. Fiebach, H.C. Friederich, C. Stippich, M. Weisbrod, S. Kaiser, Neural reward processing is modulated by approach- and avoidance-related personality traits, NeuroImage 49(2) (2010) 1868–74.

[37] P.R. Goldin, K. McRae, W. Ramel, J.J. Gross, The neural bases of emotion regulation: reappraisal and suppression of negative emotion, Biological psychiatry 63(6) (2008) 577–86.

[38] G.J. Quirk, J.S. Beer, Prefrontal involvement in the regulation of emotion: convergence of rat and human studies, Current opinion in neurobiology 16(6) (2006) 723–7.

[39] A. Etkin, T. Egner, R. Kalisch, Emotional processing in anterior cingulate and medial prefrontal cortex, Trends in cognitive sciences 15(2) (2011) 85–93.

[40] K.N. Ochsner, J.J. Gross, The cognitive control of emotion, Trends in cognitive sciences 9(5) (2005) 242–9.

[41] C.D. Salzman, S. Fusi, Emotion, cognition, and mental state representation in amygdala and prefrontal cortex, Annual review of neuroscience 33 (2010) 173–202.

[42] F.A. Mansouri, M.J. Buckley, K. Tanaka, The essential role of primate orbitofrontal cortex in conflict-induced executive control adjustment, The Journal of neuroscience: the official journal of the Society for Neuroscience 34(33) (2014) 11016–31.

[43] E.C. Cieslik, K. Zilles, S. Caspers, C. Roski, T.S. Kellermann, O. Jakobs, R. Langner, A.R. Laird, P.T. Fox, S.B. Eickhoff, Is there “one” DLPFC in cognitive action control? Evidence for heterogeneity from co-activation-based parcellation, Cerebral cortex 23(11) (2013) 2677–89.

[44] R.L. Aupperle, A.J. Melrose, A. Francisco, M.P. Paulus, M.B. Stein, Neural substrates of approach-avoidance conflict decision-making, Human brain mapping 36(2) (2015) 449–62.

[45] E.G. Chrysikou, C. Gorey, R.L. Aupperle, Anodal transcranial direct current stimulation over right dorsolateral prefrontal cortex alters decision making during approach-avoidance conflict, Social cognitive and affective neuroscience 12(3) (2017) 468–475.

[46] M.M. Botvinick, J.D. Cohen, C.S. Carter, Conflict monitoring and anterior cingulate cortex: an update, Trends in cognitive sciences 8(12) (2004) 539–46.

[47] C.S. Carter, V. van Veen, Anterior cingulate cortex and conflict detection: an update of theory and data, Cognitive, affective & behavioral neuroscience 7(4) (2007) 367–79.

[48] K. Amemori, A.M. Graybiel, Localized microstimulation of primate pregenual cingulate cortex induces negative decision-making, Nature neuroscience 15(5) (2012) 776–85.

[49] M. Hoistad, H. Barbas, Sequence of information processing for emotions through pathways linking temporal and insular cortices with the amygdala, NeuroImage 40(3) (2008) 1016–33.

[50] J.T. Buhle, J.A. Silvers, T.D. Wager, R. Lopez, C. Onyemekwu, H. Kober, J. Weber, K.N. Ochsner, Cognitive reappraisal of emotion: a meta-analysis of human neuroimaging studies, Cerebral cortex 24(11) (2014) 2981–90.

[51] S.F. White, C. Adalio, Z.T. Nolan, J. Yang, A. Martin, J.R. Blair, The amygdala’s response to face and emotional information and potential category-specific modulation of temporal cortex as a function of emotion, Frontiers in human neuroscience 8 (2014) 714.

[52] J.F. Cryan, F.F. Sweeney, The age of anxiety: role of animal models of anxiolytic action in drug discovery, British journal of pharmacology 164(4) (2011) 1129–61.

[53] M.J. Millan, The neurobiology and control of anxious states, Progress in neurobiology 70(2) (2003) 83–244.

[54] J. Song, C. Davey, C. Poulsen, P. Luu, S. Turovets, E. Anderson, K. Li, D. Tucker, EEG source localization: Sensor density and head surface coverage, Journal of neuroscience methods 256 (2015) 9–21.

[55] K. Wendel, O. Vaisanen, J. Malmivuo, N.G. Gencer, B. Vanrumste, P. Durka, R. Magjarevic, S. Supek, M.L. Pascu, H. Fontenelle, R. Grave de Peralta Menendez, EEG/MEG source imaging: methods, challenges, and open issues, Computational intelligence and neuroscience (2009) 656092.

[56] O. Hauk, Keep it simple: a case for using classical minimum norm estimation in the analysis of EEG and MEG data, NeuroImage 21(4) (2004) 1612–21.

