## Supplementary figures and images for "A translational approach to measuring the neural systems underlying approach-avoidance conflict in humans"

### Supplemental Figure 1

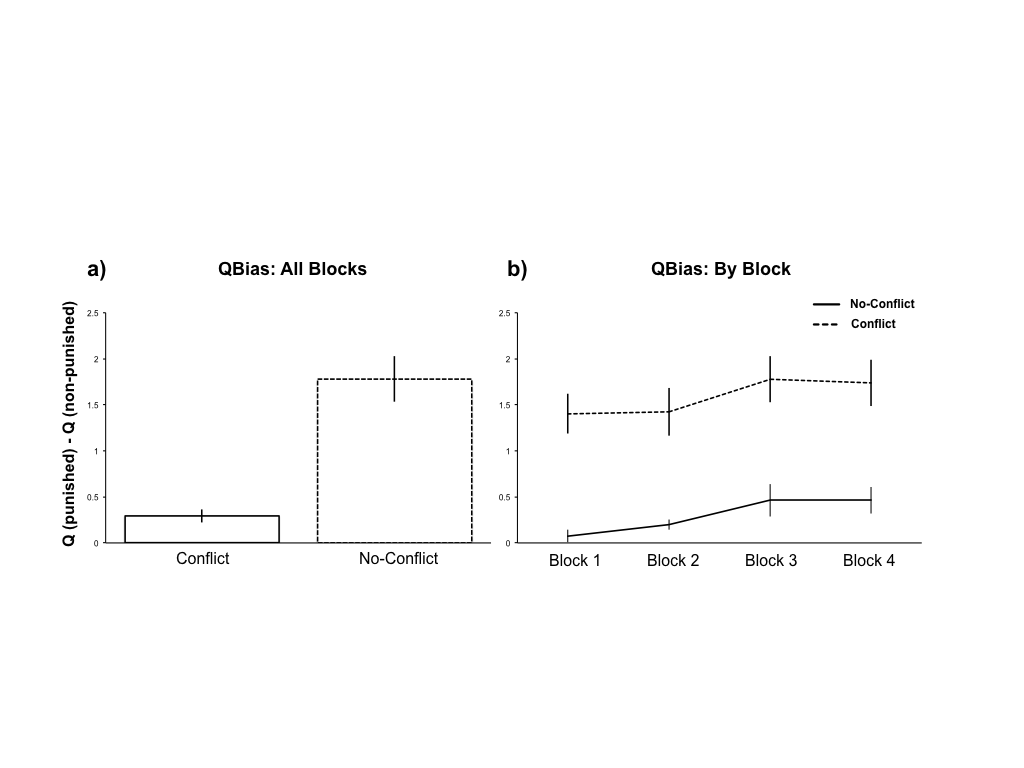
