## Supplemental Materials for "A translational approach to measuring the neural systems underlying approach-avoidance conflict in humans"

**Supplemental Methods and Materials**

*Task Design*

**Task Parameters.** Juice was delivered using a serial-triggered peristaltic syringe pump (<http://www.syringepump.com>). Shock was delivered using a parallel (LPT)-triggered Grass**^®^** SD-9 Stimulator. The juice was reinforced randomly, and independently on each side at a 10% reinforcement schedule (1/10 responses), which remained constant throughout each block. The shock was presented mid-way through the block, associated with responses on one side of the screen only. The shock was presented at a 5% reinforcement rate, which increased every 3 shocks by .05%. The reinforcement rate of the shock is increased such that behavior can be modeled as a function of the punishment rate to provide additional variables for enhanced sensitivity in behavioral analyses. Reward was held constant to provide a solid benchmark to compare punishment against. A given task session is composed of four blocks, with shocked-side randomized for each block. The shock could range from 1-10V in intensity, and the juice was delivered in .3mL-drop rewards for each participant.

**Experimental Conditions**. All task assessments were performed with the participant isolated from the research staff, such that a curtain separated the experimenters from the participants to ensure that participant behavior and cognition was not influenced by social factors resulting from the presence of the experimenter in the room (Winkel and Sarason, 1964; Ingle, 1974; Huguet et al., 1999; Belletier et al., 2015; Belletier and Camos, 2018; Chapman et al., 2018). Participants were monitored to ensure experimental compliance through video streaming from a concealed, rear-mounted webcam (Logitech® HD Pro Webcam C920) to a computer monitor mounted on the experimenter-side of the room.

**Shock Workup.** To standardize the aversiveness of the punishment across participants, we embedded a shock workup in the procedure, in order to ensure that the in-task punishment resulted in equivalent discomfort across individuals. The shock workup was automated to ensure participants were not biased by the social influence of reporting their evaluations to an experimenter (Winkel and Sarason, 1964; Ingle, 1974; Huguet et al., 1999; Belletier et al., 2015; Belletier and Camos, 2018; Chapman et al., 2018). Therefore, the entire workup procedure was conducted with the participant at the computer, and the experimenter adjusting the shock intensity (as this cannot be automated) while sitting behind and out of line of sight of participant. The voltage adjustments were based on masked codes outputted to the experimenter by an internal algorithm. During thresholding, participants were asked to report their discomfort after each shock by pressing a number on the keyboard representing a scale from 1 to 9, with 1 being “barely noticeable”, 5 being “very uncomfortable”, and 9 being “absolutely intolerable”. An automated algorithm would begin the thresholding at 1V, moving up in increments of 1V until the participant reported a discomfort level of 3, or the experimenter reached 3V, whichever came first. From there, the algorithm would move in .25V increments, increasing with ratings below 5, and decreasing with ratings above 5, until a voltage eliciting a rating of 5 was reported. At this point, the shock intensity was locked to this value.

**Juice Workup.** To begin, participants were surveyed on their juice preference prior to their study visit. This information was used to determine which juice they would receive during the task. Additionally, participants were asked to refrain from eating and drinking for 3 hours prior to the experiment. Compliance was assessed by surveying the reported times of last meal and drink. Finally, prior to each task block, participants were administered a single oyster cracker (which dried out their mouth) to provide further motivation for juice.

**In-Task Shock-Juice Equilibration.** Following the workup, participants entered a thresholding block in which they were instructed that for the next 30 seconds, they could use the up and down arrow to gain a juice reward, where the up arrow resulted in juice more often, but sometimes resulted in shock, while the down arrow resulted in less juice and was not paired with shock. In this block, the number of consecutive shocks were increased with up arrow presses until the participant stopped using the up arrow. This could result in a threshold of 1-5 consecutive shocks, at the initial thresholded intensity. This provided an in-task thresholding, directly pairing juice and shock against each other, to further ensure that the intensity of the shock was in fact calibrated in value to the juice.

**In-Task Shock-Juice Assessment.** As an assessment of the effect of the shock thresholding, a second block preceded the task in which participants were told to use a button(s) to move an object from the bottom to the top of the screen. They were instructed that the up arrow would result in juice and sometimes shock, and that the down arrow would result in neither juice nor shock. Importantly, they were instructed that both arrows would get the object to the top of the screen at the same rate. This assessment block allowed the experimental, post-hoc analyses of the efficacy of the thresholding, as the utility of the up arrow in this context would only be driven by the motivation for juice. Therefore, if the juice and shock were calibrated effectively, we would expect participants not to disproportionately use the down arrow, as this would suggest the value of the shock outweighed the value of the juice.

**Action Value.** Based on an econometrics modeling, we assumed that the choice probabilities could be characterized by the difference between the action values as: *P*_right_ (*t*) = 1/(1 + exp(−(*Q*_right_ (*t*) – *Q*_left_ (*t*)))), where *Q*_right_ (*t*) and *Q*_left_ (*t*) are the action value function of the right and left target selection (Train, 2009). The outcome value in trial *t* is modelled as *v*(*t*) = *r*(*t*) + *κ* * *pun*(*t*). We set *r*(*t*) = 1 for reward and *r*(*t*) = 0 for no reward, *pun*(*t*) = 1 for punishment, and *pun*(*t*) = 0 for no punishment. The parameter *κ* thus denoted the cost-benefit ratio of reward and punishment (Amemori and Graybiel, 2012). The action values for the chosen option *i* are updated as follows: *Q_i_*(*t* + 1) = *Q_i_*(*t*) + *α* (*v*(*t*) − *Qi*(*t*)), where α is the learning rate, where the reward prediction error was denoted by, (t) = *v*(*t*) − *Q_i_*(*t*). We performed optimization procedure using a matlab function *fminsearch* to derive the parameter *κ* and *α*, that could predict the choice pattern.

*EEG Data Processing*

All EEG data analyses were performed in Matlab (R2014b, The Mathworks Inc., MA) using custom scripts built upon the preprocessing packages in EEGLAB (Delorme and Makeig, 2004) and ARTIST (Wu et al., 2018) toolboxes. Following import into Matlab, the data was cleaned using the following pipeline 1) Data was notch filtered to remove 60 Hz line noise and its harmonic, and bandpass filtered between 1 Hz and 55 Hz using a zero-phase finite impulse response filter 2) Bad channels were rejected by computing the maximum correlation coefficient of the EEG at each electrode with the rest of the electrodes on the continuous EEG data. Bad electrodes were identified based on spatial correlations and interpolated from the EEG of adjacent channels (Wu et al., 2018). Subjects with more than 20% bad channels were discarded. 3) Remaining artifacts (ocular, heartbeat, and high-frequency persistent muscle artifact) were removed using Independent Component Analyses (ICA). ICA was applied to the data to generate spatio-temporal-spectral decomposition of the signal, and non-electrophysiological artifacts were identified in the independent components and removed using the ARTIST algorithm (Wu et al., 2018). Independent components related to the scalp muscle artifact, ocular artifact, ECG artifact, were automatically rejected using a pattern classifier trained on expert-labeled ICs from another independent EEG data set (Wu et al., 2018); 4) EEG data were re-referenced to the common average. 5) Finally, data were binned into 10-second time bins, concatenated across blocks.

Source localization to the cortical surface was performed using custom script. A three-layer symmetric boundary element model of the head was computed based on a Montreal Neurological Institute (MNI) brain template (regularization parameter = 0.1, depth-weighting component=0.5, noise-covariance estimation procedure = identity matrix). Rotating dipoles at 3003 vertices were generated on the cortical surface. The lead-field matrix was obtained by projecting the standard electrode positions onto the scalp. For each subject, an imaging kernel that maps from the channel space EEG to the source space current density was then estimated by the minimum norm estimation approach with depth weighting and regularization. For each vertex, the current density time series were reduced from their 3 orthogonal axes to a single principal direction by PCA.

*Statistical Analyses*

Voxels found to associate with behavior differentially depending on task condition were then grouped into clusters, with boundaries defined by spatial contiguity. Those clusters whose maximal voxel-to-voxel Euclidian distance was greater than 5% of the total Euclidian distance across the cortex were divided into spatially-defined subclusters using k-means. The number of subclusters were defined as the ratio of the cluster maximal voxel-to-voxel Euclidian distance to the squared cortical maximal voxel-to-voxel Euclidian distance. Finally, the vertice with peak activation was extracted from each cluster as representative of the greater cluster, and the Brodmann area (Brodmann, 1909) within which it fell was identified.

**Supplemental Results**

**Shock Workup.** Participants were thresholded on average to 4.49V +/- 1.39V. There was no effect of gender with this thresholding procedure (Male: 4.58 +/- 1.39; Female: 4.50 +/- 1.45; T(37)=-0.18, p=0.86).

**Shock and Juice “Likability” Ratings.** In-task ratings revealed that the task elicited motivationally-evocative outcomes across participants, with shock and juice both eliciting significant non-zero outcome likability ratings (Figure 2b; Shock = 4.78 +/- 1.26, two-sided one-sample t-test T(27)=12.60, p<.001; Juice = 4.34 +/- 1.82, one-sample T(27)=19.99, p<.001). The task additionally elicited motivationally balanced outcomes across participants, with juice and shock eliciting statistically equivalent absolute likability ratings (Figure 2b; two-sided paired t-test T(27)=-1.08, p=.29), and 3) outcomes with reliable value across blocks (Figure 2b; ICC: Juice = .96 , Shock=.85). Post-task ratings, in which subjects rated both the likability of shock and juice on the same scale (-10 to 10), further revealed the statistically balanced value of juice and shock across participants (Figure 2a; Shock = -4.44 +/- 2.32; Juice = 4.74 +/- 3.08, two-sided paired t-test T(39)=-.51, p=.61). Lastly, participants were asked to rate how motivated they felt by the shock and juice independently, on a scale from 1 to 10. Participants reported a statistically equivalent motivation to both shock and juice (Shock = 6.46 +/- 2.72, Juice = 6.79 +/- 2.66; two-sided paired t-test T(36)=-.08, p=0.93).

**In-Task Shock-Juice Equilibration.** Participants responded with the up arrow during this block an average of 70.2% +/- 35.6% of the time.

**In-Task Shock-Juice Assessment.** Participants utilized the up arrow (resulting in juice and shock) to achieve the goal of this block 60.5% +/- 33.7% of the time, suggesting a variable but effective thresholding which led participants, on average, to approach juice reward despite the associated shock punishment approximately 60% of the time.

**Q_Bias_.** After correcting for multiple comparisons across all 12012 (3003 vertices, 4 spectral bands), 181 distinct clusters were found to associate with bin-to-bin variability in in-task avoidance behavior differentially by task condition, as measured by Q_Bias_ (see Table S3; Figure 7a). As with response bias, the majority of these clusters were within frontal and parietal cortices, with the rest falling within the visual, somatosensory/motor, temporal, or cingulate cortices. The effects were found globally across all frequency bands.

We next sought to understand which of these significant EEG features were driven by behavior-EEG associations specific to the conflict condition (Figure 7b). We ran mixed models for the two task conditions separately, further constraining analyses to only those EEG features identified as significant in the full analysis using a p-value threshold of 0.05. Greater power predicted less Q_Bias_ behavior during conflict trials within frontal and visual areas, whereas greater power predicted greater Q_Bias_ behavior during conflict trials within temporal and motor areas. While the frontal and visual effects were found across frequency bands, the temporal effects were isolated within the alpha and theta bands.

**Supplemental Discussion**

*Validation of a Novel, Translational Approach-Avoidance Conflict Task in Humans*

Animal research has made great progress by employing targeted assays of approach-avoidance conflict to identify neural systems underlying this behavior (Kirlic et al., 2017). As with many fields, animal research allows for the experimental control in hypothesis testing that is either not ethical or simply not possible in human work. However, the mapping of rodent neuroanatomy and behavior to humans remains unclear (Gosling, 2001; Hackam and Redelmeier, 2006; Berridge and Kringelbach, 2008). Therefore, the translation of animal work to humans is critical for the elucidation of clinically-relevant neural substrates, and eventual development of novel therapeutic approaches. While there are a number of factors that have made it difficult to generalize from non-human to human work, one prominent factor has been the limitations of existing task paradigms designed to study this behavior in humans.

One way to bridge the animal and human literature is by developing human analogues to currently-used animal paradigms (Kirlic et al., 2017). One primary challenge to effectively doing this lies in the complexity of human behavior, as relevant, but confounding contextual components such as risk and reward/punishment value significantly impact decision-making, inducing uncontrolled variability in behavioral estimates. Given the inherent heterogeneity of behavioral responding in approach-avoidance, another challenge is the current lack of individualized thresholding in tasks to ensure each person is (1) motivated sufficiently by the outcomes embedded in the task and (2) motivated as much by the punishing as the rewarding outcomes. Therefore, the first step to identifying the neural substrates driving approach-avoidance conflict behavior, at its core, requires the operationalization of a paradigm stripped of confounding elements, and implementing punishing and rewarding outcomes that are both evocative and equivalent in their associated value.

The current paradigm is the first human paradigm, to our knowledge, to utilize primary reinforcers (shock, juice) for both reward and punishment. While a number of studies have implemented primary reinforcers for in-task punishment (Talmi et al., 2009; Schlund et al., 2016; Kirlic et al., 2017), none have done so simultaneously for the reward. Therefore, such paradigms are inherently more susceptible to between-subject behavioral variability resultant from imbalanced reward and punishment value, making approach- and avoidance-motivated decisions difficult to disentangle from task-specific factors (i.e., how much a subject cares about gaining points).

Importantly, the current paradigm implements a series of thresholdings to calibrate the intensity of punishment with the reward to minimize the confounding effects of imbalanced outcomes. Further, assessments of outcome value during and after the task allows for its inclusion as covariates in data analyses. The validation of the proposed paradigm suggests that 1) the thresholding effectively produces balanced and evocative reward and punishment, and 2) participant behavior elicited in this task is relatively stable.

As such, the current task provides a powerful translational bridge to animal work. The EEG features found in this study to associate with bin-to-bin variability in behavior, differentially by task condition, closely reflect those most relevant to conflict-specific approach- and avoidance- drives.

*Limitations*

The current experiment provides a step forward in measuring approach-avoidance behavior in humans. The study draws attention to the need to validate tasks by using thresholding, reliability assessments, and controlled settings. Like any task, this task also has limitations. One limitation is that participants received variable amounts of shock, which could drive distinct behavioral trajectories within the task. Therefore, controlling for this factor is critical in interpretation of results. Further, the juice is not calibrated in intensity, and therefore the value of the juice can only be driven by the pre-experimental factors (i.e, cracker administration, food/water deprivation). Another critical limitation is that to ensure the amount of juice was equilibrated across subjects, the task was designed to be self-paced. However, this results in variable task duration, possibly resulting in variable levels of task-fatigue across participants.

**Supplemental Figures.**

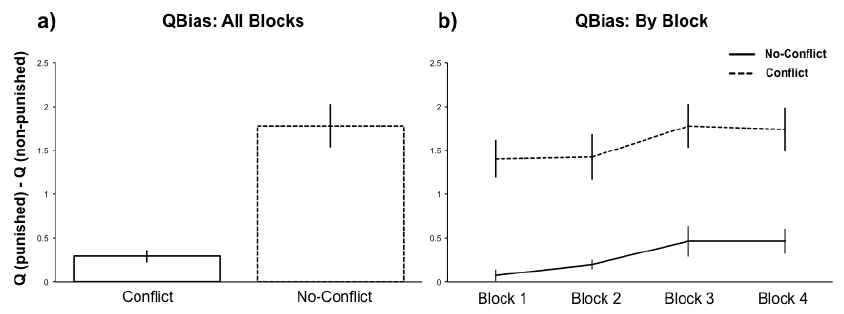

**Figure S1**. Assessment of in-task Q_Bias_: a) The action values of the punished response was greater than non-punished blocks specific to conflict blocks (T(37)=-6.16, p<.001). b) Stability of in-task behavior across blocks was assessed by ICC (Conflict = .95, No-Conflict = .53).

**Supplemental Tables.**

**Supplemental Table S1.** Statistically significant spectral condition-dependent EEG features.

| **Band** | **z** | **p** | **Vertex** | **X** | **Y** | **Z** | **BA*** | **Hem*** | **Region** (*peak activation*) |
| --- | --- | --- | --- | --- | --- | --- | --- | --- | --- |
| Theta | -2.21 | 0.05 | 1326 | -6.08 | -1.83 | 83.67 | 23 | R | cingulate cortex |
| Theta | 5.69 | 0.00 | 240 | -50.13 | -45.64 | 74.23 | 19 | R | associative visual cortex (V3, V4 & V5) |
| Theta | 3.50 | 0.00 | 288 | -47.78 | -48.00 | 53.28 | 19 | R | associative visual cortex (V3, V4 & V5) |
| Theta | -2.20 | 0.05 | 1069 | -14.93 | -26.07 | 32.28 | 37 | R | fusiform gyrus |
| Theta | -2.21 | 0.05 | 2334 | 43.52 | 26.98 | 19.24 | 11 | L | orbitofrontal area |
| Theta | -2.22 | 0.05 | 2090 | 33.85 | 39.43 | 86.89 | 9 | L | dorsolateral prefrontal cortex |
| Theta | 6.15 | 0.00 | 390 | -42.16 | 50.03 | 74.30 | 39 | L | angular gyrus |
| Theta | 8.15 | 0.00 | 574 | -36.67 | 41.44 | 99.09 | 7 | L | superior parietal lobule |
| Theta | 12.34 | 0.00 | 652 | -29.22 | 51.12 | 93.17 | 7 | L | superior parietal lobule |
| Theta | 7.17 | 0.00 | 2958 | 88.62 | 14.54 | 27.16 | 11 | L | orbitofrontal area |
| Theta | 2.43 | 0.03 | 212 | -52.70 | 44.71 | 74.74 | 19 | L | associative visual cortex (V3, V4 & V5) |
| Theta | -2.24 | 0.05 | 319 | -36.49 | -28.37 | 78.75 | 19 | R | associative visual cortex (V3, V4 & V5) |
| Theta | 7.19 | 0.00 | 487 | -35.23 | -44.08 | 97.05 | 7 | R | superior parietal lobule |
| Theta | 4.72 | 0.00 | 637 | -27.35 | -58.19 | 60.09 | 39 | R | angular gyrus |
| Theta | 6.45 | 0.00 | 776 | -21.56 | -6.27 | 119.3  8 | 5 | R | superior parietal lobule |
| Theta | 8.23 | 0.00 | 547 | -34.29 | -16.62 | 105.1  8 | 7 | R | superior parietal lobule |
| Theta | 3.81 | 0.00 | 343 | -45.97 | 49.38 | 64.64 | 39 | L | angular gyrus |
| Theta | 7.40 | 0.00 | 1668 | 15.74 | 43.76 | 109.3  2 | 4 | L | primary motor cortex |
| Theta | 8.71 | 0.00 | 793 | -20.96 | 40.51 | 105.2  8 | 5 | L | superior parietal lobule |
| Theta | 6.14 | 0.00 | 796 | -23.63 | 6.30 | 116.3  0 | 5 | L | superior parietal lobule |
| Theta | -2.22 | 0.05 | 427 | -36.29 | 24.29 | 81.29 | 19 | L | associative visual cortex (V3, V4 & V5) |
| Theta | 5.53 | 0.00 | 1156 | -10.73 | -67.05 | 63.31 | 22 | R | superior temporal gyrus |
| Theta | 6.28 | 0.00 | 1287 | -2.70 | -56.94 | 97.54 | 40 | R | supramarginal gyrus |
| Theta | 8.25 | 0.00 | 2815 | 76.57 | -12.96 | 18.30 | 11 | R | orbitofrontal area |
| Theta | 10.05 | 0.00 | 1854 | 17.77 | -49.06 | 3.26 | 20 | R | inferior temporal gyrus |
| Theta | 6.69 | 0.00 | 1692 | 13.31 | -49.74 | 105.5 | 1 | R | primary somatosensory cortex |

|  |  |  |  |  |  | 0 |  |  |  |
| --- | --- | --- | --- | --- | --- | --- | --- | --- | --- |
| Theta | 2.88 | 0.01 | 1779 | 19.13 | -64.98 | 75.84 | 2 | R | primary somatosensory cortex |
| Theta | 9.65 | 0.00 | 1963 | 24.17 | -43.94 | 2.66 | 20 | R | inferior temporal gyrus |
| Theta | -2.31 | 0.04 | 1217 | 0.47 | 34.28 | 24.67 | 20 | L | inferior temporal gyrus |
| Theta | 3.46 | 0.00 | 1585 | 11.16 | 66.31 | 29.06 | 22 | L | superior temporal gyrus |
| Theta | -2.56 | 0.02 | 2030 | 30.30 | 56.81 | 27.74 | 22 | L | superior temporal gyrus |
| Theta | -2.27 | 0.04 | 1893 | 22.12 | 27.93 | 8.11 | 20 | L | inferior temporal gyrus |
| Theta | -2.21 | 0.05 | 1393 | 0.70 | 5.77 | 96.35 | 4 | L | primary motor cortex |
| Theta | -2.21 | 0.05 | 1514 | 8.16 | -6.18 | 96.13 | 4 | R | primary motor cortex |
| Theta | 5.77 | 0.00 | 1988 | 25.37 | -47.68 | 100.7  9 | 6 | R | premotor cortex  and supplementary motor cortex |
| Theta | 6.07 | 0.00 | 2594 | 59.74 | -34.74 | 91.40 | 9 | R | dorsolateral prefrontal cortex |
| Theta | -2.40 | 0.03 | 2076 | 29.01 | 42.47 | 9.65 | 21 | L | middle temporal gyrus |
| Theta | 4.24 | 0.00 | 2070 | 28.47 | 22.27 | 115.6  4 | 6 | L | premotor cortex  and supplementary motor cortex |
| Theta | 2.83 | 0.01 | 2183 | 33.49 | 35.85 | 104.9  4 | 6 | L | premotor cortex  and supplementary motor cortex |
| Theta | 3.49 | 0.00 | 2496 | 54.98 | 47.77 | 30.89 | 11 | L | orbitofrontal area |
| Theta | -4.43 | 0.00 | 2537 | 60.39 | -3.24 | 99.90 | 8 | R | frontal eye fields |
| Theta | -2.42 | 0.03 | 3002 | 96.49 | -11.62 | 45.41 | 10 | R | anterior prefrontal cortex |
| Theta | -2.50 | 0.03 | 2696 | 70.22 | -42.26 | 47.29 | 46 | R | dorsolateral prefrontal cortex |
| Theta | 2.23 | 0.05 | 2592 | 62.53 | 4.80 | 34.13 | 24 | L | cingulate cortex |
| Theta | -2.24 | 0.05 | 2709 | 68.36 | 12.53 | 60.38 | 32 | L | dorsal anterior cingulate area |
| Theta | -2.34 | 0.04 | 2847 | 74.67 | -6.24 | 65.78 | 9 | R | dorsolateral prefrontal cortex |
| Theta | 4.51 | 0.00 | 2996 | 95.54 | 8.60 | 57.02 | 10 | L | anterior prefrontal cortex |
| Alpha | 8.79 | 0.00 | 1963 | 24.17 | -43.94 | 2.66 | 20 | R | inferior temporal gyrus |
| Alpha | 6.23 | 0.00 | 136 | -58.37 | -37.48 | 66.54 | 19 | R | associative visual cortex (V3, V4  & V5) |
| Alpha | 9.29 | 0.00 | 1854 | 17.77 | -49.06 | 3.26 | 20 | R | inferior temporal gyrus |
| Alpha | 8.49 | 0.00 | 421 | -42.21 | -16.47 | 103.7  7 | 7 | R | superior parietal lobule |
| Alpha | -2.26 | 0.04 | 1147 | -6.30 | -42.47 | 63.11 | 42 | R | primary auditory cortex |
| Alpha | 8.26 | 0.00 | 646 | -29.52 | -24.94 | 114.5  1 | 7 | R | superior parietal lobule |
| Alpha | -2.37 | 0.03 | 93 | -63.58 | 23.75 | 36.35 | 18 | L | secondary visual cortex (V2) |
| Alpha | 7.46 | 0.00 | 2955 | 88.64 | 5.81 | 23.36 | 11 | L | orbitofrontal area |
| Alpha | 9.20 | 0.00 | 623 | -29.60 | 52.76 | 88.99 | 7 | L | superior parietal lobule |
| Alpha | 3.67 | 0.00 | 1585 | 11.16 | 66.31 | 29.06 | 22 | L | superior temporal gyrus |
| Alpha | 12.01 | 0.00 | 652 | -29.22 | 51.12 | 93.17 | 7 | L | superior parietal lobule |
| Alpha | 5.78 | 0.00 | 1895 | 21.95 | 53.39 | 97.44 | 4 | L | primary motor cortex |
| Alpha | -2.48 | 0.03 | 95 | -62.78 | 32.46 | 45.98 | 18 | L | secondary visual cortex (V2) |
| Alpha | -2.30 | 0.04 | 359 | -43.39 | 25.54 | 81.64 | 19 | L | associative visual cortex (V3, V4 |

|  |  |  |  |  |  |  |  |  | & V5) |
| --- | --- | --- | --- | --- | --- | --- | --- | --- | --- |
| Alpha | 3.15 | 0.00 | 188 | -57.52 | -15.05 | 92.05 | 19 | R | associative visual cortex (V3, V4  & V5) |
| Alpha | 2.44 | 0.03 | 212 | -52.70 | 44.71 | 74.74 | 19 | L | associative visual cortex (V3, V4  & V5) |
| Alpha | 3.65 | 0.00 | 265 | -49.06 | 50.32 | 62.82 | 19 | L | associative visual cortex (V3, V4  & V5) |
| Alpha | 2.80 | 0.01 | 451 | -37.86 | 54.97 | 60.52 | 39 | L | angular gyrus |
| Alpha | 5.88 | 0.00 | 1287 | -2.70 | -56.94 | 97.54 | 40 | R | supramarginal gyrus |
| Alpha | 4.90 | 0.00 | 1156 | -10.73 | -67.05 | 63.31 | 22 | R | superior temporal gyrus |
| Alpha | 6.35 | 0.00 | 2594 | 59.74 | -34.74 | 91.40 | 9 | R | dorsolateral prefrontal cortex |
| Alpha | 4.25 | 0.00 | 1774 | 16.00 | -42.22 | 107.8  9 | 4 | R | primary motor cortex |
| Alpha | 6.16 | 0.00 | 1988 | 25.37 | -47.68 | 100.7  9 | 6 | R | premotor cortex  and supplementary motor cortex |
| Alpha | 5.62 | 0.00 | 2070 | 28.47 | 22.27 | 115.6  4 | 6 | L | premotor cortex  and supplementary motor cortex |
| Alpha | -2.22 | 0.05 | 2663 | 61.40 | 8.43 | 102.9  8 | 8 | L | frontal eye fields |
| Alpha | 2.98 | 0.01 | 1804 | 17.99 | -32.81 | 19.68 | 36 | R | perirhinal cortex & ectorhinal  area |
| Alpha | 3.08 | 0.01 | 2183 | 33.49 | 35.85 | 104.9  4 | 6 | L | premotor cortex  and supplementary motor cortex |
| Alpha | -2.26 | 0.04 | 2546 | 56.02 | -6.05 | 91.40 | 8 | R | frontal eye fields |
| Alpha | -2.30 | 0.04 | 2655 | 62.64 | -43.61 | 53.81 | 46 | R | dorsolateral prefrontal cortex |
| Alpha | -2.27 | 0.04 | 3002 | 96.49 | -11.62 | 45.41 | 10 | R | anterior prefrontal cortex |
| Alpha | -2.22 | 0.05 | 2795 | 77.17 | -5.28 | 58.13 | 9 | R | dorsolateral prefrontal cortex |
| Alpha | 2.79 | 0.01 | 2811 | 77.95 | 20.84 | 80.82 | 9 | L | dorsolateral prefrontal cortex |
| Beta | 7.02 | 0.00 | 599 | -35.79 | -51.50 | 74.01 | 39 | R | angular gyrus |
| Beta | 5.24 | 0.00 | 164 | -58.65 | -34.13 | 66.52 | 19 | R | associative visual cortex (V3, V4  & V5) |
| Beta | 8.79 | 0.00 | 421 | -42.21 | -16.47 | 103.7  7 | 7 | R | superior parietal lobule |
| Beta | -2.22 | 0.05 | 522 | -36.37 | 2.49 | 68.84 | 23 | L | cingulate cortex |
| Beta | 7.52 | 0.00 | 793 | -20.96 | 40.51 | 105.2  8 | 5 | L | superior parietal lobule |
| Beta | 6.12 | 0.00 | 1741 | 18.24 | 42.59 | 111.7  7 | 4 | L | primary motor cortex |
| Beta | -2.30 | 0.04 | 241 | -50.55 | 42.36 | 61.10 | 19 | L | associative visual cortex (V3, V4  & V5) |
| Beta | -2.33 | 0.04 | 174 | -56.43 | 26.26 | 64.81 | 19 | L | associative visual cortex (V3, V4  & V5) |

| Beta | 10.53 | 0.00 | 790 | -23.07 | 53.18 | 94.83 | 40 | L | supramarginal gyrus |
| --- | --- | --- | --- | --- | --- | --- | --- | --- | --- |
| Beta | 11.64 | 0.00 | 652 | -29.22 | 51.12 | 93.17 | 7 | L | superior parietal lobule |
| Beta | 4.23 | 0.00 | 343 | -45.97 | 49.38 | 64.64 | 39 | L | angular gyrus |
| Beta | 3.16 | 0.00 | 451 | -37.86 | 54.97 | 60.52 | 39 | L | angular gyrus |
| Beta | -2.41 | 0.03 | 910 | -16.24 | 58.72 | 35.34 | 21 | L | middle temporal gyrus |
| Beta | -2.44 | 0.03 | 903 | -19.35 | 51.74 | 54.25 | 22 | L | superior temporal gyrus |
| Beta | 5.95 | 0.00 | 2175 | 32.41 | -45.42 | 4.90 | 21 | R | middle temporal gyrus |
| Beta | -2.23 | 0.05 | 1060 | -12.78 | -46.54 | 49.04 | 22 | R | superior temporal gyrus |
| Beta | 5.63 | 0.00 | 1287 | -2.70 | -56.94 | 97.54 | 40 | R | supramarginal gyrus |
| Beta | 7.75 | 0.00 | 1854 | 17.77 | -49.06 | 3.26 | 20 | R | inferior temporal gyrus |
| Beta | -2.42 | 0.03 | 1046 | -10.35 | 59.71 | 65.19 | 42 | L | primary auditory cortex |
| Beta | 6.45 | 0.00 | 1692 | 13.31 | -49.74 | 105.5  0 | 1 | R | primary somatosensory cortex |
| Beta | 5.96 | 0.00 | 1365 | -1.38 | -46.19 | 109.1  3 | 1 | R | primary somatosensory cortex |
| Beta | -2.26 | 0.04 | 1304 | 1.24 | -19.60 | 103.9  4 | 4 | R | primary motor cortex |
| Beta | 4.26 | 0.00 | 1585 | 11.16 | 66.31 | 29.06 | 22 | L | superior temporal gyrus |
| Beta | 3.31 | 0.00 | 1800 | 21.73 | 59.66 | 15.81 | 21 | L | middle temporal gyrus |
| Beta | -2.45 | 0.03 | 1508 | 7.71 | -63.04 | 65.68 | 43 | R | primary gustatory cortex |
| Beta | 5.39 | 0.00 | 1988 | 25.37 | -47.68 | 100.7  9 | 6 | R | premotor cortex  and supplementary motor cortex |
| Beta | 3.02 | 0.01 | 1556 | 11.08 | -40.01 | 111.7  4 | 4 | R | primary motor cortex |
| Beta | 5.96 | 0.00 | 2594 | 59.74 | -34.74 | 91.40 | 9 | R | dorsolateral prefrontal cortex |
| Beta | -2.53 | 0.02 | 2504 | 54.62 | -53.58 | 45.21 | 47 | R | pars orbitalis, part of the inferior  frontal gyrus |
| Beta | -2.26 | 0.04 | 2047 | 33.51 | -62.67 | 62.46 | 4 | R | primary motor cortex |
| Beta | 7.36 | 0.00 | 2815 | 76.57 | -12.96 | 18.30 | 11 | R | orbitofrontal area |
| Beta | 5.88 | 0.00 | 2629 | 62.79 | -20.90 | 24.40 | 11 | R | orbitofrontal area |
| Beta | 3.15 | 0.00 | 2451 | 52.32 | -54.00 | 67.04 | 45 | R | Brocas area |
| Beta | -2.32 | 0.04 | 2427 | 48.92 | 36.29 | 23.47 | 11 | L | orbitofrontal area |
| Beta | 4.98 | 0.00 | 2996 | 95.54 | 8.60 | 57.02 | 10 | L | anterior prefrontal cortex |
| Beta | 4.55 | 0.00 | 2908 | 84.24 | 36.80 | 35.91 | 11 | L | orbitofrontal area |
| Beta | 8.28 | 0.00 | 2943 | 87.07 | 12.36 | 23.46 | 11 | L | orbitofrontal area |
| Beta | -2.31 | 0.04 | 2546 | 56.02 | -6.05 | 91.40 | 8 | R | frontal eye fields |
| Beta | 2.32 | 0.04 | 2553 | 62.15 | -1.91 | 37.74 | 24 | R | cingulate cortex |
| Beta | -2.43 | 0.03 | 2864 | 78.75 | -15.84 | 77.88 | 9 | R | dorsolateral prefrontal cortex |
| Beta | -2.25 | 0.04 | 2897 | 82.68 | -37.62 | 55.34 | 10 | R | anterior prefrontal cortex |
| Gamma | 8.43 | 0.00 | 547 | -34.29 | -16.62 | 105.1  8 | 7 | R | superior parietal lobule |
| Gamma | 2.61 | 0.02 | 1804 | 17.99 | -32.81 | 19.68 | 36 | R | perirhinal cortex & ectorhinal |

|  |  |  |  |  |  |  |  |  | area |
| --- | --- | --- | --- | --- | --- | --- | --- | --- | --- |
| Gamma | 4.82 | 0.00 | 164 | -58.65 | -34.13 | 66.52 | 19 | R | associative visual cortex (V3, V4  & V5) |
| Gamma | 7.12 | 0.00 | 599 | -35.79 | -51.50 | 74.01 | 39 | R | angular gyrus |
| Gamma | 4.05 | 0.00 | 1585 | 11.16 | 66.31 | 29.06 | 22 | L | superior temporal gyrus |
| Gamma | -2.24 | 0.05 | 151 | -54.88 | 46.58 | 36.74 | 19 | L | associative visual cortex (V3, V4  & V5) |
| Gamma | -2.23 | 0.05 | 315 | -46.71 | 10.90 | 61.47 | 17 | L | primary visual cortex (V1) |
| Gamma | -2.31 | 0.04 | 1869 | 20.44 | 19.92 | 11.01 | 20 | L | inferior temporal gyrus |
| Gamma | 2.20 | 0.05 | 1800 | 21.73 | 59.66 | 15.81 | 21 | L | middle temporal gyrus |
| Gamma | -2.38 | 0.03 | 95 | -62.78 | 32.46 | 45.98 | 18 | L | secondary visual cortex (V2) |
| Gamma | 4.42 | 0.00 | 343 | -45.97 | 49.38 | 64.64 | 39 | L | angular gyrus |
| Gamma | 10.24 | 0.00 | 790 | -23.07 | 53.18 | 94.83 | 40 | L | supramarginal gyrus |
| Gamma | 11.55 | 0.00 | 652 | -29.22 | 51.12 | 93.17 | 7 | L | superior parietal lobule |
| Gamma | 6.19 | 0.00 | 824 | -21.07 | 4.70 | 112.4  6 | 5 | L | superior parietal lobule |
| Gamma | 5.57 | 0.00 | 414 | -39.01 | 17.86 | 110.7  0 | 7 | L | superior parietal lobule |
| Gamma | 7.44 | 0.00 | 793 | -20.96 | 40.51 | 105.2  8 | 5 | L | superior parietal lobule |
| Gamma | 2.91 | 0.01 | 451 | -37.86 | 54.97 | 60.52 | 39 | L | angular gyrus |
| Gamma | 2.75 | 0.01 | 1311 | 2.57 | -57.79 | 24.83 | 21 | R | middle temporal gyrus |
| Gamma | 5.88 | 0.00 | 1287 | -2.70 | -56.94 | 97.54 | 40 | R | supramarginal gyrus |
| Gamma | 8.76 | 0.00 | 1854 | 17.77 | -49.06 | 3.26 | 20 | R | inferior temporal gyrus |
| Gamma | 7.99 | 0.00 | 2815 | 76.57 | -12.96 | 18.30 | 11 | R | orbitofrontal area |
| Gamma | 4.88 | 0.00 | 2552 | 55.38 | -30.27 | 24.05 | 11 | R | orbitofrontal area |
| Gamma | -2.33 | 0.04 | 1046 | -10.35 | 59.71 | 65.19 | 42 | L | primary auditory cortex |
| Gamma | -2.22 | 0.05 | 1201 | -5.66 | 64.65 | 73.89 | 40 | L | supramarginal gyrus |
| Gamma | 5.75 | 0.00 | 1365 | -1.38 | -46.19 | 109.1  3 | 1 | R | primary somatosensory cortex |
| Gamma | 6.44 | 0.00 | 1692 | 13.31 | -49.74 | 105.5  0 | 1 | R | primary somatosensory cortex |
| Gamma | -2.22 | 0.05 | 1508 | 7.71 | -63.04 | 65.68 | 43 | R | primary gustatory cortex |
| Gamma | 5.89 | 0.00 | 1741 | 18.24 | 42.59 | 111.7  7 | 4 | L | primary motor cortex |
| Gamma | 5.05 | 0.00 | 2070 | 28.47 | 22.27 | 115.6  4 | 6 | L | premotor cortex  and supplementary motor cortex |
| Gamma | -2.41 | 0.03 | 1460 | 4.35 | -20.72 | 109.8  1 | 4 | R | primary motor cortex |
| Gamma | 2.64 | 0.02 | 2661 | 63.63 | -21.46 | 89.89 | 8 | R | frontal eye fields |
| Gamma | 5.96 | 0.00 | 2594 | 59.74 | -34.74 | 91.40 | 9 | R | dorsolateral prefrontal cortex |
| Gamma | 5.20 | 0.00 | 1988 | 25.37 | -47.68 | 100.7 | 6 | R | premotor cortex |

|  |  |  |  |  |  | 9 |  |  | and supplementary motor cortex |
| --- | --- | --- | --- | --- | --- | --- | --- | --- | --- |
| Gamma | -2.23 | 0.05 | 2094 | 26.96 | -23.64 | 100.4  4 | 6 | R | premotor cortex  and supplementary motor cortex |
| Gamma | 0.00 | 0.04 | 2062 | 32.01 | -38.20 | 49.96 |  |  | #N/A |
| Gamma | 0.00 | 0.05 | 1663 | 13.16 | -41.25 | 47.23 |  |  | #N/A |
| Gamma | -2.27 | 0.04 | 2354 | 46.40 | -50.80 | 56.64 | 45 | R | Brocas area |
| Gamma | -2.49 | 0.03 | 1677 | 14.77 | -64.92 | 60.30 | 43 | R | primary gustatory cortex |
| Gamma | -2.21 | 0.05 | 2436 | 49.78 | 45.32 | 84.12 | 9 | L | dorsolateral prefrontal cortex |
| Gamma | -2.34 | 0.04 | 2791 | 69.17 | 14.43 | 94.80 | 8 | L | frontal eye fields |
| Gamma | 2.64 | 0.02 | 2451 | 52.32 | -54.00 | 67.04 | 45 | R | Brocas area |
| Gamma | 3.50 | 0.00 | 2496 | 54.98 | 47.77 | 30.89 | 11 | L | orbitofrontal area |
| Gamma | 4.80 | 0.00 | 2974 | 91.09 | 19.90 | 37.77 | 10 | L | anterior prefrontal cortex |
| Gamma | 4.64 | 0.00 | 2908 | 84.24 | 36.80 | 35.91 | 11 | L | orbitofrontal area |
| Gamma | 8.10 | 0.00 | 2943 | 87.07 | 12.36 | 23.46 | 11 | L | orbitofrontal area |
| Gamma | 2.46 | 0.03 | 2722 | 68.99 | 2.37 | 39.25 | 24 | L | cingulate cortex |
| Gamma | -2.36 | 0.03 | 2864 | 78.75 | -15.84 | 77.88 | 9 | R | dorsolateral prefrontal cortex |
| Gamma | -2.49 | 0.03 | 2988 | 94.64 | -13.37 | 56.79 | 10 | R | anterior prefrontal cortex |
| Gamma | 2.29 | 0.04 | 2811 | 77.95 | 20.84 | 80.82 | 9 | L | dorsolateral prefrontal cortex |

*BA = Brodmann Area; Hem = Hemisphere

**Supplemental Table S2.** Cortical band power differentiating behavior, measured by response bias, by task condition.

| **Band** | **z** | **p** | **Vertex** | **X** | **Y** | **Z** | **BA*** | **Hem*** | **Region** (*peak activation*) |
| --- | --- | --- | --- | --- | --- | --- | --- | --- | --- |
| Theta | 5.55 | 0.00 | 563 | -35.55 | 7.46 | 48.49 | 18 | L | secondary visual cortex (V2) |
| Theta | 6.31 | 0.00 | 523 | -33.38 | 42.69 | 42.34 | 37 | L | fusiform gyrus |
| Theta | 7.19 | 0.00 | 113 | -64.23 | 9.93 | 49.68 | 17 | L | primary visual cortex (V1) |
| Theta | 8.21 | 0.00 | 2824 | 75.46 | -10.98 | 17.22 | 11 | R | orbitofrontal area |
| Theta | 8.21 | 0.00 | 278 | -49.34 | -18.15 | 85.09 | 19 | R | associative visual cortex (V3, V4 & V5) |
| Theta | 5.08 | 0.00 | 643 | -35.49 | -5.14 | 49.03 | 18 | R | secondary visual cortex (V2) |
| Theta | 7.13 | 0.00 | 2255 | 45.07 | -3.32 | 20.97 | 25 | R | subgenual area |
| Theta | 7.99 | 0.00 | 2929 | 85.37 | -25.19 | 66.93 | 9 | R | dorsolateral prefrontal cortex |
| Theta | 8.13 | 0.00 | 1311 | 2.57 | -57.79 | 24.83 | 21 | R | middle temporal gyrus |
| Theta | 2.99 | 0.01 | 205 | -54.26 | 24.72 | 66.06 | 19 | L | associative visual cortex (V3, V4 & V5) |
| Theta | 6.60 | 0.00 | 518 | -35.33 | -50.49 | 80.47 | 39 | R | angular gyrus |
| Theta | 6.24 | 0.00 | 1802 | 18.90 | -23.41 | 107.83 | 4 | R | primary motor cortex |
| Theta | 5.20 | 0.00 | 1460 | 4.35 | -20.72 | 109.81 | 4 | R | primary motor cortex |
| Theta | 5.22 | 0.00 | 679 | -28.34 | -22.74 | 103.27 | 7 | R | superior parietal lobule |
| Theta | 7.64 | 0.00 | 483 | -36.92 | 49.64 | 87.16 | 39 | L | angular gyrus |
| Theta | 7.12 | 0.00 | 389 | -39.90 | 43.23 | 90.65 | 7 | L | superior parietal lobule |
| Theta | 2.38 | 0.04 | 537 | -34.68 | 26.38 | 99.39 | 7 | L | superior parietal lobule |
| Theta | 3.02 | 0.01 | 580 | -32.65 | -60.52 | 55.64 | 39 | R | angular gyrus |
| Theta | 3.60 | 0.00 | 1361 | 0.60 | 36.27 | 107.08 | 3 | L | primary somatosensory cortex |
| Theta | 3.64 | 0.00 | 964 | -14.13 | 6.40 | 116.84 | 1 | L | primary somatosensory cortex |
| Theta | 4.97 | 0.00 | 712 | -25.13 | 60.74 | 77.42 | 40 | L | supramarginal gyrus |
| Theta | 6.04 | 0.00 | 1441 | 3.08 | -61.47 | 90.38 | 40 | R | supramarginal gyrus |
| Theta | 2.80 | 0.01 | 973 | -10.19 | 49.65 | 45.15 | 22 | L | superior temporal gyrus |
| Theta | 4.42 | 0.00 | 1115 | -7.66 | 33.16 | 104.26 | 2 | L | primary somatosensory cortex |
| Theta | 7.18 | 0.00 | 2541 | 48.47 | 2.13 | 25.41 | 32 | L | dorsal anterior cingulate area |
| Theta | 8.21 | 0.00 | 2582 | 56.55 | 55.28 | 49.36 | 45 | L | broca area |
| Theta | 8.21 | 0.00 | 2716 | 68.86 | 4.51 | 30.38 | 32 | L | dorsal anterior cingulate area |
| Theta | 5.08 | 0.00 | 1892 | 24.29 | 57.39 | 52.17 | 43 | L | primary gustatory cortex |
| Theta | 7.40 | 0.00 | 2016 | 27.17 | 56.42 | 12.83 | 21 | L | middle temporal gyrus |
| Theta | 4.69 | 0.00 | 1079 | -10.85 | -66.27 | 59.84 | 22 | R | superior temporal gyrus |
| Theta | 2.67 | 0.02 | 1064 | -9.65 | -49.75 | 45.60 | 22 | R | superior temporal gyrus |
| Theta | 2.39 | 0.04 | 1437 | 4.65 | 3.02 | 91.72 | 31 | L | dorsal posterior cingulate area |
| Theta | 4.97 | 0.00 | 1716 | 15.05 | 3.92 | 108.04 | 4 | L | primary motor cortex |
| Theta | 4.39 | 0.00 | 2023 | 28.40 | 2.70 | 105.61 | 6 | L | premotor cortex  and supplementary motor cortex |

| Theta | 2.35 | 0.04 | 1643 | 13.57 | -3.44 | 90.11 | 24 | R | cingulate cortex |
| --- | --- | --- | --- | --- | --- | --- | --- | --- | --- |
| Theta | 3.78 | 0.00 | 1704 | 15.75 | -62.91 | 77.51 | 2 | R | primary somatosensory cortex |
| Theta | 3.99 | 0.00 | 1745 | 16.79 | -60.22 | 87.15 | 1 | R | primary somatosensory cortex |
| Theta | 4.70 | 0.00 | 2032 | 33.78 | 47.84 | 95.22 | 6 | L | premotor cortex  and supplementary motor cortex |
| Theta | 2.26 | 0.05 | 1724 | 12.96 | 52.36 | 32.83 | 22 | L | superior temporal gyrus |
| Theta | 2.25 | 0.05 | 1820 | 20.70 | -53.73 | 55.13 | 43 | R | primary gustatory cortex |
| Theta | 2.47 | 0.03 | 1954 | 25.44 | -54.00 | 95.73 | 6 | R | premotor cortex  and supplementary motor cortex |
| Theta | 7.88 | 0.00 | 2958 | 88.62 | 14.54 | 27.16 | 11 | L | orbitofrontal area |
| Theta | 7.11 | 0.00 | 2965 | 89.59 | -6.05 | 28.84 | 11 | R | orbitofrontal area |
| Alpha | 7.07 | 0.00 | 730 | -24.77 | 30.37 | 41.26 | 37 | L | fusiform gyrus |
| Alpha | 8.13 | 0.00 | 623 | -29.60 | 52.76 | 88.99 | 7 | L | superior parietal lobule |
| Alpha | 8.01 | 0.00 | 2716 | 68.86 | 4.51 | 30.38 | 32 | L | dorsal anterior cingulate area |
| Alpha | 7.41 | 0.00 | 94 | -64.62 | 13.44 | 51.26 | 17 | L | primary visual cortex (V1) |
| Alpha | 8.13 | 0.00 | 2574 | 60.50 | 52.36 | 54.64 | 45 | L | broca area |
| Alpha | 4.98 | 0.00 | 1716 | 15.05 | 3.92 | 108.04 | 4 | L | primary motor cortex |
| Alpha | 7.76 | 0.00 | 1498 | 10.59 | -56.81 | 15.77 | 21 | R | middle temporal gyrus |
| Alpha | 7.76 | 0.00 | 237 | -53.02 | -15.19 | 88.23 | 19 | R | associative visual cortex (V3, V4  & V5) |
| Alpha | 6.61 | 0.00 | 1802 | 18.90 | -23.41 | 107.83 | 4 | R | primary motor cortex |
| Alpha | 8.13 | 0.00 | 2649 | 71.13 | -9.50 | 21.87 | 11 | R | orbitofrontal area |
| Alpha | 7.97 | 0.00 | 21 | -68.88 | -13.73 | 53.75 | 17 | R | primary visual cortex (V1) |
| Alpha | 2.64 | 0.02 | 419 | -39.21 | -41.09 | 50.65 | 19 | R | associative visual cortex (V3, V4  & V5) |
| Alpha | 2.80 | 0.01 | 534 | -34.61 | -40.51 | 59.87 | 39 | R | angular gyrus |
| Alpha | 3.09 | 0.01 | 663 | -20.32 | -44.11 | 33.53 | 37 | R | fusiform gyrus |
| Alpha | 2.58 | 0.02 | 691 | -26.48 | 36.12 | 108.91 | 7 | L | superior parietal lobule |
| Alpha | 5.25 | 0.00 | 712 | -25.13 | 60.74 | 77.42 | 40 | L | supramarginal gyrus |
| Alpha | 5.09 | 0.00 | 1115 | -7.66 | 33.16 | 104.26 | 2 | L | primary somatosensory cortex |
| Alpha | 3.17 | 0.00 | 1506 | 6.03 | 54.69 | 81.08 | 2 | L | primary somatosensory cortex |
| Alpha | 4.42 | 0.00 | 1156 | -10.73 | -67.05 | 63.31 | 22 | R | superior temporal gyrus |
| Alpha | 2.36 | 0.04 | 1024 | -14.08 | -56.05 | 31.40 | 21 | R | middle temporal gyrus |
| Alpha | 2.52 | 0.03 | 1464 | 5.87 | 67.09 | 52.16 | 22 | L | superior temporal gyrus |
| Alpha | 3.71 | 0.00 | 1704 | 15.75 | -62.91 | 77.51 | 2 | R | primary somatosensory cortex |
| Alpha | 2.48 | 0.03 | 1820 | 20.70 | -53.73 | 55.13 | 43 | R | primary gustatory cortex |
| Alpha | 2.70 | 0.02 | 1724 | 12.96 | 52.36 | 32.83 | 22 | L | superior temporal gyrus |
| Alpha | 2.33 | 0.04 | 2729 | 69.87 | -12.87 | 57.21 | 32 | R | dorsal anterior cingulate area |
| Alpha | 2.34 | 0.04 | 2737 | 68.55 | 26.05 | 64.92 | 9 | L | dorsolateral prefrontal cortex |
| Alpha | 3.04 | 0.01 | 2857 | 80.62 | -9.38 | 88.93 | 9 | R | dorsolateral prefrontal cortex |
| Alpha | 7.93 | 0.00 | 2976 | 91.69 | 7.20 | 30.72 | 10 | L | anterior prefrontal cortex |
| Alpha | 2.87 | 0.01 | 2949 | 88.15 | 23.14 | 50.83 | 10 | L | anterior prefrontal cortex |

| Alpha | 7.82 | 0.00 | 2965 | 89.59 | -6.05 | 28.84 | 11 | R | orbitofrontal area |
| --- | --- | --- | --- | --- | --- | --- | --- | --- | --- |
| Alpha | 2.30 | 0.04 | 2970 | 90.21 | 9.38 | 72.12 | 9 | L | dorsolateral prefrontal cortex |
| Beta | 7.99 | 0.00 | 389 | -39.90 | 43.23 | 90.65 | 7 | L | superior parietal lobule |
| Beta | 6.17 | 0.00 | 635 | -26.02 | 27.33 | 42.25 | 37 | L | fusiform gyrus |
| Beta | 6.23 | 0.00 | 94 | -64.62 | 13.44 | 51.26 | 17 | L | primary visual cortex (V1) |
| Beta | 6.72 | 0.00 | 523 | -33.38 | 42.69 | 42.34 | 37 | L | fusiform gyrus |
| Beta | 7.94 | 0.00 | 2962 | 90.94 | -24.41 | 58.19 | 10 | R | anterior prefrontal cortex |
| Beta | 8.21 | 0.00 | 1614 | 5.83 | -57.30 | 23.47 | 21 | R | middle temporal gyrus |
| Beta | 7.44 | 0.00 | 278 | -49.34 | -18.15 | 85.09 | 19 | R | associative visual cortex (V3, V4  & V5) |
| Beta | 4.05 | 0.00 | 643 | -35.49 | -5.14 | 49.03 | 18 | R | secondary visual cortex (V2) |
| Beta | 6.00 | 0.00 | 21 | -68.88 | -13.73 | 53.75 | 17 | R | primary visual cortex (V1) |
| Beta | 8.13 | 0.00 | 2789 | 78.35 | -5.08 | 33.19 | 10 | R | anterior prefrontal cortex |
| Beta | 5.13 | 0.00 | 1460 | 4.35 | -20.72 | 109.81 | 4 | R | primary motor cortex |
| Beta | 6.22 | 0.00 | 1842 | 18.01 | -20.35 | 109.86 | 4 | R | primary motor cortex |
| Beta | 7.48 | 0.00 | 518 | -35.33 | -50.49 | 80.47 | 39 | R | angular gyrus |
| Beta | 7.03 | 0.00 | 646 | -29.52 | -24.94 | 114.51 | 7 | R | superior parietal lobule |
| Beta | 4.02 | 0.00 | 580 | -32.65 | -60.52 | 55.64 | 39 | R | angular gyrus |
| Beta | 2.90 | 0.01 | 903 | -19.35 | 51.74 | 54.25 | 22 | L | superior temporal gyrus |
| Beta | 3.86 | 0.00 | 973 | -10.19 | 49.65 | 45.15 | 22 | L | superior temporal gyrus |
| Beta | 2.29 | 0.04 | 691 | -26.48 | 36.12 | 108.91 | 7 | L | superior parietal lobule |
| Beta | 5.07 | 0.00 | 712 | -25.13 | 60.74 | 77.42 | 40 | L | supramarginal gyrus |
| Beta | 2.39 | 0.04 | 719 | -26.20 | -53.86 | 79.56 | 40 | R | supramarginal gyrus |
| Beta | 4.48 | 0.00 | 1369 | -0.19 | -62.56 | 89.37 | 40 | R | supramarginal gyrus |
| Beta | 3.46 | 0.00 | 1361 | 0.60 | 36.27 | 107.08 | 3 | L | primary somatosensory cortex |
| Beta | 4.04 | 0.00 | 1716 | 15.05 | 3.92 | 108.04 | 4 | L | primary motor cortex |
| Beta | 3.69 | 0.00 | 1026 | -10.09 | 4.76 | 121.84 | 3 | L | primary somatosensory cortex |
| Beta | 4.44 | 0.00 | 1115 | -7.66 | 33.16 | 104.26 | 2 | L | primary somatosensory cortex |
| Beta | 8.21 | 0.00 | 2833 | 79.55 | 7.85 | 17.01 | 11 | L | orbitofrontal area |
| Beta | 4.04 | 0.00 | 1122 | -5.84 | 62.19 | 70.52 | 40 | L | supramarginal gyrus |
| Beta | 7.79 | 0.00 | 2489 | 53.36 | 51.67 | 48.94 | 45 | L | broca area |
| Beta | 6.24 | 0.00 | 2541 | 48.47 | 2.13 | 25.41 | 32 | L | dorsal anterior cingulate area |
| Beta | 7.41 | 0.00 | 2016 | 27.17 | 56.42 | 12.83 | 21 | L | middle temporal gyrus |
| Beta | 2.52 | 0.03 | 890 | -17.22 | 45.78 | 91.88 | 7 | L | superior parietal lobule |
| Beta | 3.70 | 0.00 | 1156 | -10.73 | -67.05 | 63.31 | 22 | R | superior temporal gyrus |
| Beta | 2.72 | 0.02 | 988 | -11.90 | -32.84 | 100.39 | 2 | R | primary somatosensory cortex |
| Beta | 2.34 | 0.04 | 1559 | 9.91 | -1.39 | 89.73 | 24 | R | cingulate cortex |
| Beta | 3.95 | 0.00 | 2064 | 25.91 | 52.53 | 97.68 | 6 | L | premotor cortex  and supplementary motor cortex |
| Beta | 2.48 | 0.03 | 1650 | 13.92 | 31.67 | 119.48 | 4 | L | primary motor cortex |
| Beta | 2.41 | 0.03 | 1820 | 20.70 | -53.73 | 55.13 | 43 | R | primary gustatory cortex |

| Beta | 2.28 | 0.05 | 2397 | 47.67 | -32.77 | 96.85 | 8 | R | frontal eye fields |
| --- | --- | --- | --- | --- | --- | --- | --- | --- | --- |
| Beta | 2.29 | 0.04 | 2729 | 69.87 | -12.87 | 57.21 | 32 | R | dorsal anterior cingulate area |
| Beta | 3.04 | 0.01 | 2857 | 80.62 | -9.38 | 88.93 | 9 | R | dorsolateral prefrontal cortex |
| Beta | 2.40 | 0.03 | 2822 | 75.61 | 26.88 | 78.19 | 9 | L | dorsolateral prefrontal cortex |
| Beta | 8.13 | 0.00 | 2964 | 89.57 | 7.32 | 27.72 | 11 | L | orbitofrontal area |
| Gamma | 5.87 | 0.00 | 113 | -64.23 | 9.93 | 49.68 | 17 | L | primary visual cortex (V1) |
| Gamma | 6.13 | 0.00 | 423 | -36.97 | 42.01 | 42.84 | 37 | L | fusiform gyrus |
| Gamma | 5.60 | 0.00 | 614 | -31.81 | 28.88 | 40.20 | 19 | L | associative visual cortex (V3, V4  & V5) |
| Gamma | 5.52 | 0.00 | 21 | -68.88 | -13.73 | 53.75 | 17 | R | primary visual cortex (V1) |
| Gamma | 4.29 | 0.00 | 269 | -51.82 | -2.08 | 44.55 | 18 | R | secondary visual cortex (V2) |
| Gamma | 7.21 | 0.00 | 237 | -53.02 | -15.19 | 88.23 | 19 | R | associative visual cortex (V3, V4  & V5) |
| Gamma | 4.22 | 0.00 | 200 | -52.84 | 6.08 | 84.54 | 18 | L | secondary visual cortex (V2) |
| Gamma | 4.36 | 0.00 | 365 | -52.23 | -33.68 | 84.80 | 19 | R | associative visual cortex (V3, V4  & V5) |
| Gamma | 7.90 | 0.00 | 389 | -39.90 | 43.23 | 90.65 | 7 | L | superior parietal lobule |
| Gamma | 6.74 | 0.00 | 362 | -42.83 | 39.80 | 97.28 | 7 | L | superior parietal lobule |
| Gamma | 4.52 | 0.00 | 276 | -46.83 | 10.40 | 91.61 | 7 | L | superior parietal lobule |
| Gamma | 5.34 | 0.00 | 2087 | 29.03 | -14.29 | 118.42 | 6 | R | premotor cortex  and supplementary motor cortex |
| Gamma | 6.21 | 0.00 | 679 | -28.34 | -22.74 | 103.27 | 7 | R | superior parietal lobule |
| Gamma | 7.40 | 0.00 | 518 | -35.33 | -50.49 | 80.47 | 39 | R | angular gyrus |
| Gamma | 5.49 | 0.00 | 1842 | 18.01 | -20.35 | 109.86 | 4 | R | primary motor cortex |
| Gamma | 3.51 | 0.00 | 580 | -32.65 | -60.52 | 55.64 | 39 | R | angular gyrus |
| Gamma | 2.66 | 0.02 | 688 | -27.98 | 57.32 | 68.84 | 40 | L | supramarginal gyrus |
| Gamma | 3.82 | 0.00 | 964 | -14.13 | 6.40 | 116.84 | 1 | L | primary somatosensory cortex |
| Gamma | 3.52 | 0.00 | 1781 | 15.20 | 3.82 | 116.56 | 4 | L | primary motor cortex |
| Gamma | 5.24 | 0.00 | 712 | -25.13 | 60.74 | 77.42 | 40 | L | supramarginal gyrus |
| Gamma | 3.22 | 0.00 | 719 | -26.20 | -53.86 | 79.56 | 40 | R | supramarginal gyrus |
| Gamma | 2.97 | 0.01 | 746 | -22.73 | -55.90 | 89.89 | 40 | R | supramarginal gyrus |
| Gamma | 2.83 | 0.01 | 851 | -15.69 | 29.72 | 112.47 | 2 | L | primary somatosensory cortex |
| Gamma | 3.20 | 0.00 | 919 | -14.76 | 38.25 | 106.13 | 2 | L | primary somatosensory cortex |
| Gamma | 3.99 | 0.00 | 1156 | -10.73 | -67.05 | 63.31 | 22 | R | superior temporal gyrus |
| Gamma | 3.27 | 0.00 | 1175 | -5.49 | 32.93 | 114.02 | 1 | L | primary somatosensory cortex |
| Gamma | 5.06 | 0.00 | 1441 | 3.08 | -61.47 | 90.38 | 40 | R | supramarginal gyrus |
| Gamma | 3.43 | 0.00 | 1122 | -5.84 | 62.19 | 70.52 | 40 | L | supramarginal gyrus |
| Gamma | 2.28 | 0.05 | 1141 | -6.83 | -37.74 | 109.80 | 1 | R | primary somatosensory cortex |
| Gamma | 6.56 | 0.00 | 2257 | 35.41 | -6.44 | 27.33 | 25 | R | subgenual area |
| Gamma | 8.21 | 0.00 | 2789 | 78.35 | -5.08 | 33.19 | 10 | R | anterior prefrontal cortex |
| Gamma | 5.91 | 0.00 | 2323 | 47.36 | -6.50 | 18.85 | 32 | R | dorsal anterior cingulate area |
| Gamma | 7.55 | 0.00 | 1498 | 10.59 | -56.81 | 15.77 | 21 | R | middle temporal gyrus |

| Gamma | 6.86 | 0.00 | 2016 | 27.17 | 56.42 | 12.83 | 21 | L | middle temporal gyrus |
| --- | --- | --- | --- | --- | --- | --- | --- | --- | --- |
| Gamma | 6.38 | 0.00 | 2284 | 35.85 | 3.20 | 32.66 | 25 | L | subgenual area |
| Gamma | 6.30 | 0.00 | 2006 | 27.53 | 50.63 | 10.05 | 21 | L | middle temporal gyrus |
| Gamma | 8.13 | 0.00 | 2728 | 75.32 | 1.99 | 18.60 | 11 | L | orbitofrontal area |
| Gamma | 2.71 | 0.02 | 1351 | 2.97 | -40.08 | 112.53 | 1 | R | primary somatosensory cortex |
| Gamma | 3.08 | 0.01 | 1692 | 13.31 | -49.74 | 105.50 | 1 | R | primary somatosensory cortex |
| Gamma | 2.60 | 0.02 | 1361 | 0.60 | 36.27 | 107.08 | 3 | L | primary somatosensory cortex |
| Gamma | 3.92 | 0.00 | 2064 | 25.91 | 52.53 | 97.68 | 6 | L | premotor cortex  and supplementary motor cortex |
| Gamma | 2.62 | 0.02 | 1745 | 16.79 | -60.22 | 87.15 | 1 | R | primary somatosensory cortex |
| Gamma | 6.01 | 0.00 | 2800 | 80.87 | 40.37 | 42.61 | 10 | L | anterior prefrontal cortex |
| Gamma | 6.09 | 0.00 | 2412 | 50.26 | 55.54 | 51.88 | 45 | L | broca area |
| Gamma | 7.73 | 0.00 | 2489 | 53.36 | 51.67 | 48.94 | 45 | L | broca area |
| Gamma | 3.08 | 0.01 | 2034 | 29.02 | 3.49 | 111.92 | 6 | L | premotor cortex  and supplementary motor cortex |
| Gamma | 8.13 | 0.00 | 2973 | 89.93 | -26.03 | 49.77 | 10 | R | anterior prefrontal cortex |
| Gamma | 5.34 | 0.00 | 2807 | 72.72 | -43.40 | 48.98 | 10 | R | anterior prefrontal cortex |
| Gamma | 4.07 | 0.00 | 2655 | 62.64 | -43.61 | 53.81 | 46 | R | dorsolateral prefrontal cortex |
| Gamma | 2.97 | 0.01 | 2857 | 80.62 | -9.38 | 88.93 | 9 | R | dorsolateral prefrontal cortex |
| Gamma | 6.91 | 0.00 | 2958 | 88.62 | 14.54 | 27.16 | 11 | L | orbitofrontal area |

*BA = Brodmann Area; Hem = Hemisphere

**Supplemental Table S3.** Cortical band power differentiating behavior, measured by Q_Bias_, by task condition.

| **Band** | **z** | **p** | **Vertex** | **X** | **Y** | **Z** | **BA*** | **Hem*** | **Region** (*peak activation*) |
| --- | --- | --- | --- | --- | --- | --- | --- | --- | --- |
| Theta | 5.13 | 0.00 | 146 | -58.49 | 21.79 | 71.83 | 18 | L | secondary visual cortex (V2) |
| Theta | 8.21 | 0.00 | 501 | -34.45 | 59.90 | 51.29 | 39 | L | angular gyrus |
| Theta | 8.21 | 0.00 | 126 | -61.20 | 35.76 | 59.13 | 19 | L | associative visual cortex (V3, V4 & V5) |
| Theta | 3.48 | 0.00 | 510 | -35.64 | -13.80 | 44.06 | 19 | R | associative visual cortex (V3, V4 & V5) |
| Theta | 4.14 | 0.00 | 269 | -51.82 | -2.08 | 44.55 | 18 | R | secondary visual cortex (V2) |
| Theta | 4.22 | 0.00 | 321 | -44.21 | 8.71 | 45.76 | 18 | L | secondary visual cortex (V2) |
| Theta | 3.94 | 0.00 | 802 | -18.87 | 15.11 | 39.84 | 27 | L | piriform cortex |
| Theta | 4.18 | 0.00 | 264 | -49.10 | 6.39 | 47.07 | 18 | L | secondary visual cortex (V2) |
| Theta | 8.21 | 0.00 | 2248 | 38.75 | -52.86 | 74.57 | 6 | R | premotor cortex  and supplementary motor cortex |
| Theta | 8.21 | 0.00 | 612 | -32.71 | -40.59 | 98.21 | 7 | R | superior parietal lobule |
| Theta | 8.21 | 0.00 | 1970 | 23.99 | -56.56 | 88.33 | 4 | R | primary motor cortex |
| Theta | 7.42 | 0.00 | 2754 | 70.52 | -40.15 | 44.67 | 10 | R | anterior prefrontal cortex |
| Theta | 6.31 | 0.00 | 111 | -61.04 | -14.83 | 82.53 | 18 | R | secondary visual cortex (V2) |
| Theta | 8.21 | 0.00 | 2314 | 42.56 | -41.58 | 86.47 | 9 | R | dorsolateral prefrontal cortex |
| Theta | 2.66 | 0.02 | 420 | -40.16 | 15.88 | 55.07 | 17 | L | primary visual cortex (V1) |
| Theta | 4.52 | 0.00 | 772 | -22.67 | 51.64 | 96.80 | 7 | L | superior parietal lobule |
| Theta | 6.27 | 0.00 | 723 | -25.31 | 62.02 | 64.06 | 40 | L | supramarginal gyrus |
| Theta | 6.76 | 0.00 | 1233 | -4.46 | 38.32 | 114.39 | 1 | L | primary somatosensory cortex |
| Theta | 7.58 | 0.00 | 919 | -14.76 | 38.25 | 106.13 | 2 | L | primary somatosensory cortex |
| Theta | 4.28 | 0.00 | 914 | -12.56 | -66.46 | 37.62 | 22 | R | superior temporal gyrus |
| Theta | 4.58 | 0.00 | 1815 | 20.05 | -31.97 | 1.67 | 20 | R | inferior temporal gyrus |
| Theta | 4.14 | 0.00 | 1717 | 16.95 | -39.31 | 0.95 | 20 | R | inferior temporal gyrus |
| Theta | 6.48 | 0.00 | 1562 | 9.96 | -65.20 | 70.27 | 43 | R | primary gustatory cortex |
| Theta | 8.13 | 0.00 | 1832 | 19.50 | -59.79 | 84.48 | 1 | R | primary somatosensory cortex |
| Theta | 3.44 | 0.00 | 2063 | 28.42 | -62.41 | 41.96 | 22 | R | superior temporal gyrus |
| Theta | 2.46 | 0.04 | 1035 | -10.90 | -10.22 | 109.90 | 1 | R | primary somatosensory cortex |
| Theta | 5.62 | 0.00 | 1991 | 30.98 | 21.69 | 107.81 | 6 | L | premotor cortex  and supplementary motor cortex |
| Theta | 6.05 | 0.00 | 1867 | 19.45 | 38.31 | 112.90 | 4 | L | primary motor cortex |
| Theta | 2.55 | 0.03 | 1379 | 3.89 | 55.08 | 92.61 | 2 | L | primary somatosensory cortex |
| Theta | 5.87 | 0.00 | 1876 | 22.87 | 51.65 | 24.90 | 22 | L | superior temporal gyrus |
| Theta | 5.11 | 0.00 | 2050 | 27.32 | 57.75 | 45.17 | 22 | L | superior temporal gyrus |
| Theta | 4.05 | 0.00 | 1892 | 24.29 | 57.39 | 52.17 | 43 | L | primary gustatory cortex |
| Theta | 2.40 | 0.05 | 1890 | 22.11 | 62.87 | 71.31 | 1 | L | primary somatosensory cortex |
| Theta | 3.16 | 0.01 | 2084 | 29.57 | 61.25 | 79.02 | 4 | L | primary motor cortex |

| Theta | 2.56 | 0.03 | 2183 | 33.49 | 35.85 | 104.94 | 6 | L | premotor cortex  and supplementary motor cortex |
| --- | --- | --- | --- | --- | --- | --- | --- | --- | --- |
| Theta | 3.55 | 0.00 | 2173 | 34.68 | -30.77 | 104.58 | 6 | R | premotor cortex  and supplementary motor cortex |
| Theta | 4.30 | 0.00 | 2424 | 47.95 | 57.27 | 58.70 | 44 | L | brocas area |
| Theta | 4.80 | 0.00 | 2811 | 77.95 | 20.84 | 80.82 | 9 | L | dorsolateral prefrontal cortex |
| Theta | 5.14 | 0.00 | 2822 | 75.61 | 26.88 | 78.19 | 9 | L | dorsolateral prefrontal cortex |
| Theta | 3.61 | 0.00 | 2813 | 76.49 | 4.48 | 38.02 | 32 | L | dorsal anterior cingulate area |
| Theta | 2.54 | 0.03 | 2596 | 59.87 | 43.59 | 27.29 | 11 | L | orbitofrontal area |
| Theta | 2.66 | 0.02 | 2913 | 82.60 | 21.53 | 39.34 | 10 | L | anterior prefrontal cortex |
| Theta | 2.87 | 0.01 | 2922 | 85.10 | 35.66 | 42.87 | 10 | L | anterior prefrontal cortex |
| Alpha | 6.84 | 0.00 | 89 | -64.55 | 33.11 | 63.46 | 18 | L | secondary visual cortex (V2) |
| Alpha | 7.99 | 0.00 | 610 | -32.73 | 62.26 | 55.84 | 39 | L | angular gyrus |
| Alpha | 4.59 | 0.00 | 162 | -55.28 | 3.56 | 83.64 | 18 | L | secondary visual cortex (V2) |
| Alpha | 4.16 | 0.00 | 321 | -44.21 | 8.71 | 45.76 | 18 | L | secondary visual cortex (V2) |
| Alpha | 3.73 | 0.00 | 753 | -23.73 | 11.35 | 46.17 | 19 | L | associative visual cortex (V3, V4  & V5) |
| Alpha | 4.15 | 0.00 | 264 | -49.10 | 6.39 | 47.07 | 18 | L | secondary visual cortex (V2) |
| Alpha | 4.24 | 0.00 | 269 | -51.82 | -2.08 | 44.55 | 18 | R | secondary visual cortex (V2) |
| Alpha | 3.98 | 0.00 | 301 | -46.96 | -2.09 | 50.01 | 18 | R | secondary visual cortex (V2) |
| Alpha | 8.13 | 0.00 | 2113 | 33.55 | -43.65 | 94.50 | 6 | R | premotor cortex  and supplementary motor cortex |
| Alpha | 8.01 | 0.00 | 528 | -33.90 | -32.87 | 98.21 | 7 | R | superior parietal lobule |
| Alpha | 7.13 | 0.00 | 1776 | 18.39 | -27.75 | 112.44 | 4 | R | primary motor cortex |
| Alpha | 4.24 | 0.00 | 48 | -66.47 | -28.89 | 59.48 | 18 | R | secondary visual cortex (V2) |
| Alpha | 6.51 | 0.00 | 111 | -61.04 | -14.83 | 82.53 | 18 | R | secondary visual cortex (V2) |
| Alpha | 2.80 | 0.02 | 313 | -45.06 | -48.87 | 36.99 | 37 | R | fusiform gyrus |
| Alpha | 2.57 | 0.03 | 512 | -32.89 | 42.88 | 40.19 | 37 | L | fusiform gyrus |
| Alpha | 5.01 | 0.00 | 790 | -23.07 | 53.18 | 94.83 | 40 | L | supramarginal gyrus |
| Alpha | 6.01 | 0.00 | 1876 | 22.87 | 51.65 | 24.90 | 22 | L | superior temporal gyrus |
| Alpha | 5.10 | 0.00 | 2050 | 27.32 | 57.75 | 45.17 | 22 | L | superior temporal gyrus |
| Alpha | 6.11 | 0.00 | 723 | -25.31 | 62.02 | 64.06 | 40 | L | supramarginal gyrus |
| Alpha | 7.40 | 0.00 | 1233 | -4.46 | 38.32 | 114.39 | 1 | L | primary somatosensory cortex |
| Alpha | 7.02 | 0.00 | 1305 | 0.33 | 39.91 | 116.13 | 1 | L | primary somatosensory cortex |
| Alpha | 4.51 | 0.00 | 1252 | -5.97 | -65.56 | 43.06 | 22 | R | superior temporal gyrus |
| Alpha | 4.58 | 0.00 | 1924 | 23.83 | -35.06 | -0.82 | 38 | R | temporal pole |
| Alpha | 4.98 | 0.00 | 914 | -12.56 | -66.46 | 37.62 | 22 | R | superior temporal gyrus |
| Alpha | 2.76 | 0.02 | 965 | -14.72 | -14.08 | 100.82 | 5 | R | somatosensory association  cortex (superior parietal lobule) |
| Alpha | 8.04 | 0.00 | 1560 | 11.80 | -60.08 | 76.71 | 2 | R | primary somatosensory cortex |
| Alpha | 5.16 | 0.00 | 2790 | 77.91 | -25.62 | 79.49 | 9 | R | dorsolateral prefrontal cortex |
| Alpha | 7.56 | 0.00 | 2754 | 70.52 | -40.15 | 44.67 | 10 | R | anterior prefrontal cortex |

| Alpha | 8.04 | 0.00 | 2233 | 38.52 | -59.00 | 74.56 | 6 | R | premotor cortex  and supplementary motor cortex |
| --- | --- | --- | --- | --- | --- | --- | --- | --- | --- |
| Alpha | 8.04 | 0.00 | 2434 | 50.28 | -41.40 | 85.21 | 9 | R | dorsolateral prefrontal cortex |
| Alpha | 3.02 | 0.01 | 1035 | -10.90 | -10.22 | 109.90 | 1 | R | primary somatosensory cortex |
| Alpha | 6.43 | 0.00 | 1448 | 6.67 | 31.09 | 115.07 | 4 | L | primary motor cortex |
| Alpha | 6.00 | 0.00 | 1991 | 30.98 | 21.69 | 107.81 | 6 | L | premotor cortex  and supplementary motor cortex |
| Alpha | 3.88 | 0.00 | 1892 | 24.29 | 57.39 | 52.17 | 43 | L | primary gustatory cortex |
| Alpha | 2.88 | 0.01 | 2084 | 29.57 | 61.25 | 79.02 | 4 | L | primary motor cortex |
| Alpha | 7.45 | 0.00 | 2018 | 29.02 | -52.28 | 96.58 | 6 | R | premotor cortex  and supplementary motor cortex |
| Alpha | 2.58 | 0.03 | 2061 | 32.33 | 28.07 | 100.83 | 6 | L | premotor cortex  and supplementary motor cortex |
| Alpha | 3.31 | 0.00 | 2173 | 34.68 | -30.77 | 104.58 | 6 | R | premotor cortex  and supplementary motor cortex |
| Alpha | 5.23 | 0.00 | 2717 | 68.67 | 27.57 | 77.01 | 9 | L | dorsolateral prefrontal cortex |
| Alpha | 5.51 | 0.00 | 2822 | 75.61 | 26.88 | 78.19 | 9 | L | dorsolateral prefrontal cortex |
| Alpha | 4.40 | 0.00 | 2424 | 47.95 | 57.27 | 58.70 | 44 | L | brocas area |
| Alpha | 3.18 | 0.01 | 2596 | 59.87 | 43.59 | 27.29 | 11 | L | orbitofrontal area |
| Alpha | 3.39 | 0.00 | 2928 | 85.78 | -6.93 | 39.12 | 10 | R | anterior prefrontal cortex |
| Alpha | 3.11 | 0.01 | 2813 | 76.49 | 4.48 | 38.02 | 32 | L | dorsal anterior cingulate area |
| Alpha | 2.72 | 0.02 | 2922 | 85.10 | 35.66 | 42.87 | 10 | L | anterior prefrontal cortex |
| Alpha | 2.43 | 0.04 | 2913 | 82.60 | 21.53 | 39.34 | 10 | L | anterior prefrontal cortex |
| Beta | 6.29 | 0.00 | 22 | -70.16 | 27.20 | 63.02 | 18 | L | secondary visual cortex (V2) |
| Beta | 8.21 | 0.00 | 108 | -60.15 | 42.69 | 45.22 | 19 | L | associative visual cortex (V3, V4  & V5) |
| Beta | 8.21 | 0.00 | 610 | -32.73 | 62.26 | 55.84 | 39 | L | angular gyrus |
| Beta | 8.13 | 0.00 | 682 | -24.69 | 57.25 | 42.15 | 37 | L | fusiform gyrus |
| Beta | 4.30 | 0.00 | 321 | -44.21 | 8.71 | 45.76 | 18 | L | secondary visual cortex (V2) |
| Beta | 4.05 | 0.00 | 540 | -40.95 | 14.50 | 44.55 | 18 | L | secondary visual cortex (V2) |
| Beta | 3.43 | 0.00 | 802 | -18.87 | 15.11 | 39.84 | 27 | L | piriform cortex |
| Beta | 4.48 | 0.00 | 269 | -51.82 | -2.08 | 44.55 | 18 | R | secondary visual cortex (V2) |
| Beta | 3.18 | 0.01 | 489 | -42.39 | -17.17 | 39.67 | 19 | R | associative visual cortex (V3, V4  & V5) |
| Beta | 6.17 | 0.00 | 48 | -66.47 | -28.89 | 59.48 | 18 | R | secondary visual cortex (V2) |
| Beta | 5.83 | 0.00 | 111 | -61.04 | -14.83 | 82.53 | 18 | R | secondary visual cortex (V2) |
| Beta | 7.41 | 0.00 | 2248 | 38.75 | -52.86 | 74.57 | 6 | R | premotor cortex  and supplementary motor cortex |
| Beta | 8.01 | 0.00 | 2421 | 50.04 | -43.25 | 93.13 | 8 | R | frontal eye fields |
| Beta | 8.13 | 0.00 | 528 | -33.90 | -32.87 | 98.21 | 7 | R | superior parietal lobule |
| Beta | 6.99 | 0.00 | 1776 | 18.39 | -27.75 | 112.44 | 4 | R | primary motor cortex |
| Beta | 8.21 | 0.00 | 1704 | 15.75 | -62.91 | 77.51 | 2 | R | primary somatosensory cortex |

| Beta | 7.32 | 0.00 | 2754 | 70.52 | -40.15 | 44.67 | 10 | R | anterior prefrontal cortex |
| --- | --- | --- | --- | --- | --- | --- | --- | --- | --- |
| Beta | 2.69 | 0.02 | 277 | -45.58 | 45.64 | 84.07 | 39 | L | angular gyrus |
| Beta | 2.40 | 0.05 | 420 | -40.16 | 15.88 | 55.07 | 17 | L | primary visual cortex (V1) |
| Beta | 2.57 | 0.03 | 512 | -32.89 | 42.88 | 40.19 | 37 | L | fusiform gyrus |
| Beta | 5.82 | 0.00 | 790 | -23.07 | 53.18 | 94.83 | 40 | L | supramarginal gyrus |
| Beta | 5.87 | 0.00 | 1088 | -13.04 | 65.03 | 54.12 | 22 | L | superior temporal gyrus |
| Beta | 6.66 | 0.00 | 774 | -22.64 | 64.66 | 64.63 | 40 | L | supramarginal gyrus |
| Beta | 7.79 | 0.00 | 919 | -14.76 | 38.25 | 106.13 | 2 | L | primary somatosensory cortex |
| Beta | 4.74 | 0.00 | 1265 | -3.34 | 46.84 | 104.36 | 2 | L | primary somatosensory cortex |
| Beta | 4.40 | 0.00 | 1247 | -0.52 | 60.71 | 87.92 | 40 | L | supramarginal gyrus |
| Beta | 2.40 | 0.05 | 748 | -25.30 | 12.61 | 117.13 | 5 | L | somatosensory association  cortex (superior parietal lobule) |
| Beta | 5.17 | 0.00 | 1815 | 20.05 | -31.97 | 1.67 | 20 | R | inferior temporal gyrus |
| Beta | 6.20 | 0.00 | 914 | -12.56 | -66.46 | 37.62 | 22 | R | superior temporal gyrus |
| Beta | 5.00 | 0.00 | 1924 | 23.83 | -35.06 | -0.82 | 38 | R | temporal pole |
| Beta | 2.44 | 0.04 | 1294 | -3.08 | 12.55 | 109.80 | 4 | L | primary motor cortex |
| Beta | 6.73 | 0.00 | 1448 | 6.67 | 31.09 | 115.07 | 4 | L | primary motor cortex |
| Beta | 6.68 | 0.00 | 1991 | 30.98 | 21.69 | 107.81 | 6 | L | premotor cortex  and supplementary motor cortex |
| Beta | 3.81 | 0.00 | 1872 | 25.32 | 43.09 | 27.22 | 38 | L | temporal pole |
| Beta | 5.28 | 0.00 | 1876 | 22.87 | 51.65 | 24.90 | 22 | L | superior temporal gyrus |
| Beta | 3.45 | 0.00 | 2084 | 29.57 | 61.25 | 79.02 | 4 | L | primary motor cortex |
| Beta | 2.76 | 0.02 | 2055 | 29.62 | -51.65 | 20.03 | 21 | R | middle temporal gyrus |
| Beta | 4.23 | 0.00 | 2173 | 34.68 | -30.77 | 104.58 | 6 | R | premotor cortex  and supplementary motor cortex |
| Beta | 2.76 | 0.02 | 2202 | 36.94 | 60.57 | 72.08 | 6 | L | premotor cortex  and supplementary motor cortex |
| Beta | 3.63 | 0.00 | 2367 | 45.16 | 56.57 | 59.49 | 44 | L | brocas area |
| Beta | 2.39 | 0.05 | 2327 | 43.25 | 45.11 | 87.30 | 9 | L | dorsolateral prefrontal cortex |
| Beta | 2.42 | 0.04 | 2408 | 47.43 | 56.40 | 67.17 | 44 | L | brocas area |
| Beta | 3.93 | 0.00 | 2671 | 62.91 | 50.66 | 32.13 | 47 | L | pars orbitalis, part of the inferior  frontal gyrus |
| Beta | 3.94 | 0.00 | 2650 | 64.16 | 31.06 | 82.80 | 9 | L | dorsolateral prefrontal cortex |
| Beta | 3.79 | 0.00 | 2607 | 62.83 | 32.33 | 85.19 | 9 | L | dorsolateral prefrontal cortex |
| Beta | 2.43 | 0.04 | 2722 | 68.99 | 2.37 | 39.25 | 24 | L | cingulate cortex |
| Beta | 2.44 | 0.04 | 2753 | 71.18 | 2.75 | 86.44 | 8 | L | frontal eye fields |
| Beta | 3.50 | 0.00 | 2813 | 76.49 | 4.48 | 38.02 | 32 | L | dorsal anterior cingulate area |
| Beta | 2.60 | 0.03 | 2908 | 84.24 | 36.80 | 35.91 | 11 | L | orbitofrontal area |
| Beta | 2.71 | 0.02 | 2790 | 77.91 | -25.62 | 79.49 | 9 | R | dorsolateral prefrontal cortex |
| Gamma | 8.13 | 0.00 | 108 | -60.15 | 42.69 | 45.22 | 19 | L | associative visual cortex (V3, V4  & V5) |
| Gamma | 4.16 | 0.00 | 677 | -27.60 | 18.96 | 38.78 | 19 | L | associative visual cortex (V3, V4 |

|  |  |  |  |  |  |  |  |  | & V5) |
| --- | --- | --- | --- | --- | --- | --- | --- | --- | --- |
| Gamma | 8.13 | 0.00 | 501 | -34.45 | 59.90 | 51.29 | 39 | L | angular gyrus |
| Gamma | 7.94 | 0.00 | 80 | -62.23 | 37.40 | 62.59 | 19 | L | associative visual cortex (V3, V4  & V5) |
| Gamma | 4.19 | 0.00 | 130 | -60.78 | -1.63 | 41.66 | 18 | R | secondary visual cortex (V2) |
| Gamma | 4.50 | 0.00 | 269 | -51.82 | -2.08 | 44.55 | 18 | R | secondary visual cortex (V2) |
| Gamma | 7.51 | 0.00 | 2754 | 70.52 | -40.15 | 44.67 | 10 | R | anterior prefrontal cortex |
| Gamma | 8.13 | 0.00 | 528 | -33.90 | -32.87 | 98.21 | 7 | R | superior parietal lobule |
| Gamma | 7.80 | 0.00 | 2108 | 32.02 | -51.14 | 94.08 | 6 | R | premotor cortex  and supplementary motor cortex |
| Gamma | 7.42 | 0.00 | 491 | -36.84 | -42.45 | 89.71 | 39 | R | angular gyrus |
| Gamma | 8.13 | 0.00 | 2243 | 38.13 | -57.46 | 78.01 | 4 | R | primary motor cortex |
| Gamma | 8.21 | 0.00 | 1704 | 15.75 | -62.91 | 77.51 | 2 | R | primary somatosensory cortex |
| Gamma | 2.62 | 0.03 | 420 | -40.16 | 15.88 | 55.07 | 17 | L | primary visual cortex (V1) |
| Gamma | 5.72 | 0.00 | 790 | -23.07 | 53.18 | 94.83 | 40 | L | supramarginal gyrus |
| Gamma | 6.70 | 0.00 | 843 | -15.58 | 65.59 | 57.90 | 22 | L | superior temporal gyrus |
| Gamma | 5.99 | 0.00 | 1088 | -13.04 | 65.03 | 54.12 | 22 | L | superior temporal gyrus |
| Gamma | 8.13 | 0.00 | 919 | -14.76 | 38.25 | 106.13 | 2 | L | primary somatosensory cortex |
| Gamma | 3.71 | 0.00 | 1247 | -0.52 | 60.71 | 87.92 | 40 | L | supramarginal gyrus |
| Gamma | 6.81 | 0.00 | 1483 | -0.53 | 42.73 | 114.27 | 1 | L | primary somatosensory cortex |
| Gamma | 4.57 | 0.00 | 1717 | 16.95 | -39.31 | 0.95 | 20 | R | inferior temporal gyrus |
| Gamma | 4.85 | 0.00 | 1815 | 20.05 | -31.97 | 1.67 | 20 | R | inferior temporal gyrus |
| Gamma | 6.02 | 0.00 | 914 | -12.56 | -66.46 | 37.62 | 22 | R | superior temporal gyrus |
| Gamma | 6.06 | 0.00 | 1991 | 30.98 | 21.69 | 107.81 | 6 | L | premotor cortex  and supplementary motor cortex |
| Gamma | 6.55 | 0.00 | 1867 | 19.45 | 38.31 | 112.90 | 4 | L | primary motor cortex |
| Gamma | 3.61 | 0.00 | 2081 | 33.80 | 48.59 | 33.45 | 38 | L | temporal pole |
| Gamma | 5.46 | 0.00 | 1876 | 22.87 | 51.65 | 24.90 | 22 | L | superior temporal gyrus |
| Gamma | 3.22 | 0.01 | 2084 | 29.57 | 61.25 | 79.02 | 4 | L | primary motor cortex |
| Gamma | 2.56 | 0.03 | 2055 | 29.62 | -51.65 | 20.03 | 21 | R | middle temporal gyrus |
| Gamma | 3.14 | 0.01 | 2183 | 33.49 | 35.85 | 104.94 | 6 | L | premotor cortex  and supplementary motor cortex |
| Gamma | 4.19 | 0.00 | 2173 | 34.68 | -30.77 | 104.58 | 6 | R | premotor cortex  and supplementary motor cortex |
| Gamma | 2.46 | 0.04 | 2202 | 36.94 | 60.57 | 72.08 | 6 | L | premotor cortex  and supplementary motor cortex |
| Gamma | 3.80 | 0.00 | 2367 | 45.16 | 56.57 | 59.49 | 44 | L | brocas area |
| Gamma | 2.45 | 0.04 | 2327 | 43.25 | 45.11 | 87.30 | 9 | L | dorsolateral prefrontal cortex |
| Gamma | 2.51 | 0.04 | 2448 | 50.74 | 52.12 | 66.37 | 44 | L | brocas area |
| Gamma | 3.70 | 0.00 | 2671 | 62.91 | 50.66 | 32.13 | 47 | L | pars orbitalis, part of the inferior  frontal gyrus |
| Gamma | 3.68 | 0.00 | 2650 | 64.16 | 31.06 | 82.80 | 9 | L | dorsolateral prefrontal cortex |

| Gamma | 3.98 | 0.00 | 2813 | 76.49 | 4.48 | 38.02 | 32 | L | dorsal anterior cingulate area |
| --- | --- | --- | --- | --- | --- | --- | --- | --- | --- |
| Gamma | 2.78 | 0.02 | 2913 | 82.60 | 21.53 | 39.34 | 10 | L | anterior prefrontal cortex |
| Gamma | 2.45 | 0.04 | 2753 | 71.18 | 2.75 | 86.44 | 8 | L | frontal eye fields |
| Gamma | 2.95 | 0.01 | 2790 | 77.91 | -25.62 | 79.49 | 9 | R | dorsolateral prefrontal cortex |
| Gamma | 2.38 | 0.05 | 2987 | 93.20 | 17.31 | 55.42 | 10 | L | anterior prefrontal cortex |

*BA = Brodmann Area; Hem = Hemisphere

**References**

Amemori K, Graybiel AM (2012) Localized microstimulation of primate pregenual cingulate cortex induces negative decision-making. Nature neuroscience 15:776-785.

Belletier C, Camos V (2018) Does the experimenter presence affect working memory? Annals of the New York Academy of Sciences.

Belletier C, Davranche K, Tellier IS, Dumas F, Vidal F, Hasbroucq T, Huguet P (2015) Choking under monitoring pressure: being watched by the experimenter reduces executive attention. Psychonomic bulletin & review 22:1410-1416.

Berridge KC, Kringelbach ML (2008) Affective neuroscience of pleasure: reward in humans and animals. Psychopharmacology 199:457-480.

Brodmann K (1909) Vergleichende Lokalisationslehre der Grosshirnrinde in ihren Prinzipien dargestellt auf Grund des Zellenbaues. Barth JA, Leipzig.

Chapman CD, Benedict C, Schioth HB (2018) Experimenter gender and replicability in science. Science advances 4:e1701427.

Delorme A, Makeig S (2004) EEGLAB: an open source toolbox for analysis of single-trial EEG dynamics including independent component analysis. Journal of neuroscience methods 134:9-21.

Gosling SD (2001) From mice to men: what can we learn about personality from animal research? Psychological bulletin 127:45-86.

Hackam DG, Redelmeier DA (2006) Translation of research evidence from animals to humans. Jama 296:1731-1732.

Huguet P, Galvaing MP, Monteil JM, Dumas F (1999) Social presence effects in the Stroop task: further evidence for an attentional view of social facilitation. Journal of personality and social psychology 77:1011-1025.

Ingle GM (1974) The experimenter as an audience: the effects of varied experimenter "presence" upon performance at a simple laboratory task. In. ScholarWorks: University of Massachusetts Amherst.

Kirlic N, Young J, Aupperle RL (2017) Animal to human translational paradigms relevant for approach avoidance conflict decision making. Behaviour research and therapy 96:14-29.

Schlund MW, Brewer AT, Magee SK, Richman DM, Solomon S, Ludlum M, Dymond S (2016) The tipping point: Value differences and parallel dorsal-ventral frontal circuits gating human approach-avoidance behavior. NeuroImage 136:94-105.

Talmi D, Dayan P, Kiebel SJ, Frith CD, Dolan RJ (2009) How humans integrate the prospects of pain and reward during choice. The Journal of neuroscience : the official journal of the Society for Neuroscience 29:14617-14626.

Train KE (2009) Discrete Choice Methods with Simulation: Cambridge University Press

Winkel GH, Sarason IG (1964) Subject, Experimenter, and Situational Variables in Research on Anxiety. Journal of abnormal psychology 68:601-608.

Wu W, Keller CJ, Rogasch NC, Longwell P, Shpigel E, Rolle CE, Etkin A (2018) ARTIST: A fully automated artifact rejection algorithm for single-pulse TMS-EEG data. Human brain mapping 39:1607-1625.
